# Universal alternative splicing of noncoding exons

**DOI:** 10.1101/136275

**Authors:** Ira W. Deveson, Marion E. Brunck, James Blackburn, Elizabeth Tseng, Ting Hon, Tyson A. Clark, Michael B. Clark, Joanna Crawford, Marcel E. Dinger, Lars K. Nielsen, John S. Mattick, Tim R. Mercer

## Abstract

The human transcriptome is so large, diverse and dynamic that, even after a decade of investigation by RNA sequencing (RNA-Seq), we are yet to resolve its true dimensions. RNA-Seq suffers from an expression-dependent bias that impedes characterization of low-abundance transcripts. We performed targeted single-molecule and short-read RNA-Seq to survey the transcriptional landscape of a single human chromosome (Hsa21) at unprecedented resolution. Our analysis reaches the lower limits of the transcriptome, identifying a fundamental distinction between protein-coding and noncoding gene content: almost every noncoding exon undergoes alternative splicing, producing a seemingly limitless variety of isoforms. Analysis of syntenic regions of the mouse genome shows that few noncoding exons are shared between human and mouse, yet human splicing profiles are recapitulated on Hsa21 in mouse cells, indicative of regulation by a deeply conserved splicing code. We propose that noncoding exons are functionally modular, with alternative splicing generating an enormous repertoire of potential regulatory RNAs and a rich transcriptional reservoir for gene evolution.

## INTRODUCTION

The genome is transcribed into a diverse range of protein-coding and noncoding RNAs collectively termed the transcriptome. The human transcriptome is so large and complex that, even after a decade of investigation by RNA Sequencing (RNA-Seq), we are yet to achieve a complete census of gene expression. Moreover, our view of a gene as a discrete entity, and of a single protein-coding gene as the functional unit of inheritance, has been undermined by the recognition of pervasive transcription across the genome and interleaved alternative isoforms at individual loci (Carninci et al., 2005; Clark et al., 2011; Djebali et al., 2012; Kapranov et al., 2005; Mercer et al., 2011; Sharon et al., 2013; Tilgner et al., 2015).

RNA-Seq has revealed an abundance of small and large non-protein-coding RNAs that are antisense, intronic or intergenic to protein-coding genes (Derrien et al., 2012; Hon et al., 2017; Iyer et al., 2015; You et al 2017). Similarly, many protein-coding genes express alternative isoforms that lack extended open reading frames (ORFs; Gonzàlez-Porta et al., 2013). These findings have fuelled one of the major debates of modern genetics: the functional relevance of noncoding RNA expression.

The initial sequencing of the human genome provided a catalog of around 20,000 protein-coding genes (Lander et al., 2001). However, at least as many long noncoding RNAs (lncRNAs) have since been identified, and new studies routinely discover novel genes and isoforms (Hon et al., 2017; Iyer et al., 2015; Sharon et al., 2013; Tilgner et al., 2015; You et al., 2017). This failure to achieve a comprehensive annotation of the transcriptome is partly due to the expression-dependent bias of RNA-Seq, which limits the capacity of this technique to resolve low abundance transcripts (Hardwick et al., 2016). This has impeded the discovery and characterization of lncRNAs, which are typically weakly expressed (Cabili et al., 2011; Derrien et al., 2012). As a result, our understanding of lncRNA biology has been largely informed by studying only those examples with sufficient expression for analysis by RNA-Seq.

Another limitation of traditional RNA-Seq is the reliance on computational assembly of full-length isoforms from short (~100-150 bp) sequencing reads. This is a difficult task, particularly when alternative splicing generates multiple partially redundant isoforms at an individual locus (Conesa et al., 2016). With the emergence of technologies for long-read sequencing it is now possible to read full-length isoforms as single molecules, negating the challenges posed by transcript assembly. Leading studies have highlighted the utility of single-molecule techniques for resolving complex and precisely organized alternative splicing events (Sharon et al., 2013; Tilgner et al., 2015). However, depth remains a constraint, with rare transcripts often falling below the limits of sampling.

Here, we have attempted to reach the lower limits of the transcriptome by surveying gene expression from a single human chromosome at unprecedented resolution. Chromosome 21 (Hsa21) is the smallest human chromosome (48 Mb), is typical of the human genome in many features (*e.g.* gene content and repeat density; **Figure S1**), and has accordingly been used as a model system in transcriptomics (Cawley et al., 2004; Kampa et al., 2004). With trisomy of Hsa21 being the most common chromosomal aneuploidy in liveborn children and the most frequent genetic cause of mental retardation, the gene content of this chromosome is also the subject of medical interest (Dierssen, 2012; Letourneau et al., 2014).

To resolve the complete expression profile of Hsa21, whilst excluding the remainder of the genome, we used targeted RNA sequencing (CaptureSeq; Mercer et al., 2014). Complementary oligonucleotide probes were tiled across the chromosome to capture expressed transcripts that were then sequenced deeply on single-molecule and short-read platforms. This approach reduces the influence of the expression-dependent bias inherent to RNA-Seq, allowing gene populations encoded within this cross-section of the genome to be observed at high resolution. This reveals a fundamental distinction in the architecture of protein-coding and noncoding RNA.

## RESULTS

### Targeted RNA-Seq analysis of human chromosome 21

Two limitations of traditional RNA-Seq have ensured that, despite considerable attention, the true dimensions of the human transcriptome remain unresolved. First; because sequencing reads are competitively sampled from a single pool, in which transcripts of varied abundance are proportionally represented, weakly expressed transcripts commonly evade detection (Clark et al., 2011; Hardwick et al., 2016). Second; the accurate assembly of full-length isoforms from short sequencing reads is challenging, particularly when multiple alternative isoforms are transcribed from a single locus (Conesa et al., 2016; Tilgner et al., 2015).

To address these challenges, we performed single-molecule (PacBio RSII) and short-read (Illumina HiSeq) RNA CaptureSeq, targeting the complete expression profile of Hsa21. We generated biotin-labeled oligonucleotide probes tiling the entire non-repetitive Hsa21 sequence. These were used to capture full-length cDNA molecules, thereby restricting sequencing to transcripts expressed on Hsa21 (**STAR Methods**).

To establish our approach, we first performed short-read sequencing on Hsa21-enriched cDNA libraries from the human K562 cell-type and compared these to K562 samples analyzed in parallel by conventional RNA-Seq (**STAR Methods**). Whereas just 1.3% of reads from RNA-Seq libraries were uniquely aligned to Hsa21, this figure rose to 71.8% after capture, equating to a 54-fold coverage enrichment (**Supplementary Table 1**). Analysis of ERCC spike-in controls confirmed that RNA CaptureSeq accurately measured transcript abundances within the physiological range of gene expression (**Figure S2A,B**), as previously demonstrated (Clark et al., 2015; Mercer et al., 2014). The legitimacy of novel splice-junctions identified by CaptureSeq was also verified by RT-PCR and Sanger sequencing (16/20 randomly selected examples; **Figure S2C**; **Supplementary Table 2**).

Next we analyzed Hsa21-enriched cDNA from human testis by deep single-molecule sequencing (**STAR Methods**). Testis exhibits a distinctly promiscuous transcriptional profile (Soumillon et al., 2013), marking this as an ideal tissue within which to conduct a broad survey of gene-content encoded on Hsa21. We obtained 387,029 full-length non-chimeric reads aligning to Hsa21, representing 910 Mb of usable transcript sequence concentrated in ~1.5% of the genome. After filtering, we retrieved 101,478 full-length multi-exonic transcript reads on Hsa21 (**Supplementary Table 3**; **Figure S3A,B**).

To reinforce single-molecule isoforms and, more importantly, to enable quantitative analyses of expression and splicing, we also performed very deep short-read sequencing on Hsa21-enriched cDNA from testis, brain and kidney (**STAR Methods**). An average 65-fold coverage enrichment, relative to conventional RNASeq, was achieved, and at least 100 million reads uniquely aligning to Hsa21 were obtained for each tissue (**Supplementary Table 4**). Strong correspondence between spliced short-read alignments and singlemolecule transcripts in our Hsa21 transcriptome profile was observed, with 84.7% of alignment junctions being concordant with introns in single-molecule transcripts (**Figure S3C**,**D**). Because long PacBio reads provide reliable transcript scaffolds, while read-counts for short Illumina reads accurately estimate the abundance of known isoforms, our combined CaptureSeq approach permits robust quantitative transcriptome analyses that do not rely on *de novo* transcript assembly.

### Transcriptional landscape of human chromosome 21

Hsa21 has frequently been used as a cross-sectional model for genomics because it is small (48 Mb; ~1.5% of the genome) and typical in terms of gene content, repeat density and other features (**Figure S1**; Cawley et al., 2004; Kampa et al., 2004).

Our targeted analysis showed that essentially all (non-repetitive) regions of Hsa21 encode spliced gene loci, greatly reducing intergenic regions (**Figure 1A**). This is best illustrated in two 'gene deserts' that flank the *NCAM2* gene (extending 2.6 Mb upstream and 4.0 Mb downstream), which were largely devoid of transcript annotations. We discovered that these regions harbored numerous large, multi-exonic and richly alternatively spliced lncRNAs (**Figure 1B**).

**Figure 1.**
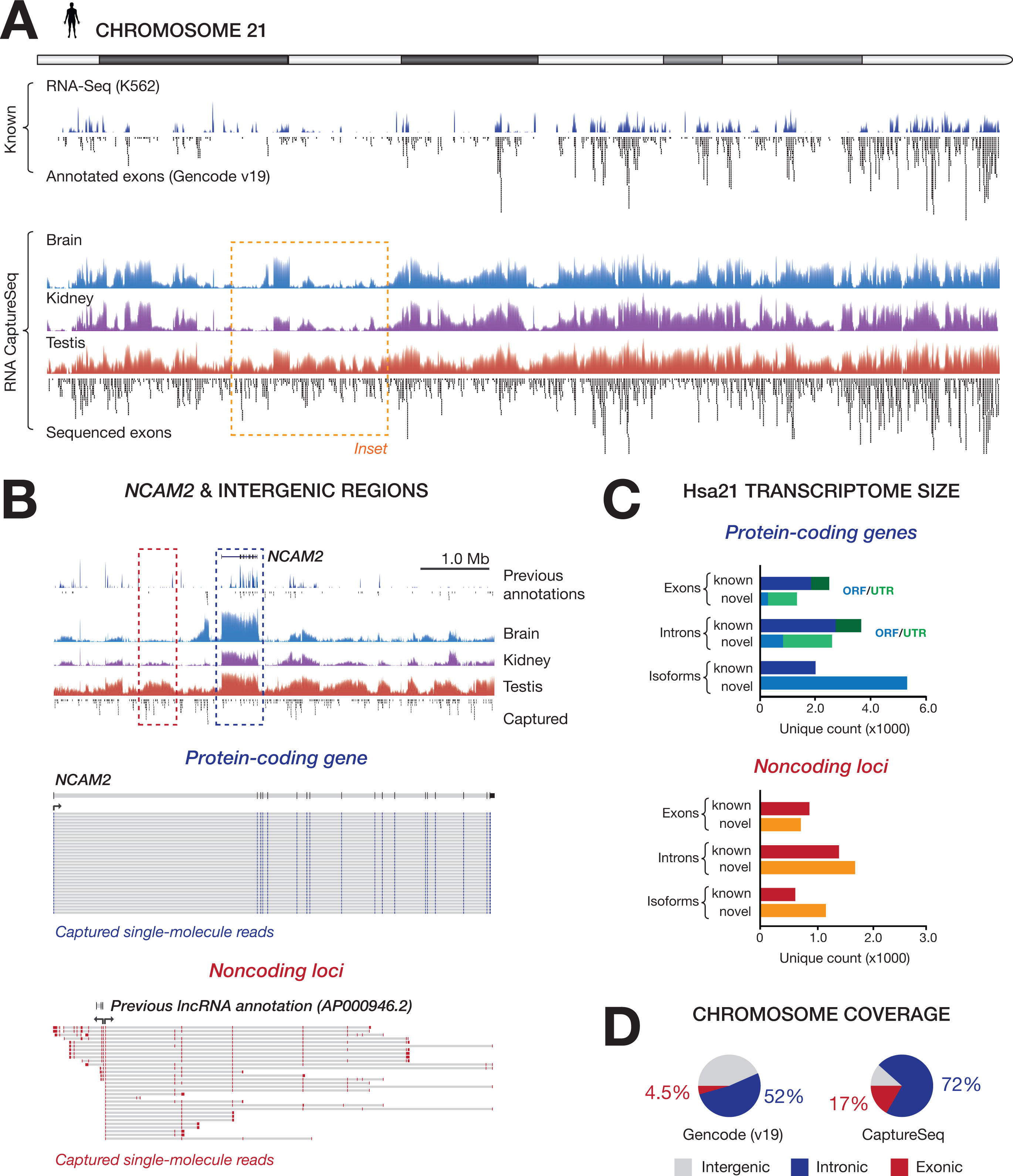
Transcriptional landscape of human chromosome 21. **(A)** Transcriptional activity recorded across the major arm of human chromosome 21 (Hsa21) by short-read and single-molecule RNA CaptureSeq. Normalized coverage (log scale) from short-read alignments is shown for brain (blue), kidney (purple) and testis (red), with coverage from traditional RNA-Seq (in K562 cells; navy) shown for comparison. Below coverage tracks are all unique internal exons (black) from transcripts resolved by single-molecule sequencing. Unique internal exons from the Gencode (v19) reference catalog are shown for comparison. **(B)** Inset from A; detail of intergenic regions flanking the protein-coding gene *NCAM2* (2.6 Mb upstream and 4.0 Mb down-stream), in which multiple novel long noncoding RNAs (lncRNAs) were detected. Red and blue insets show single-molecule reads supporting *NCAM2* (blue) and two nearby lncRNAs (red). Single-molecule RNA CaptureSeq data extends the gene model for the previously annotated lncRNA *AP000946.2* and resolves a novel lncRNA that is transcribed in the opposite direction. Both exhibit extensive alternative splicing, in contrast to *NCAM2,* where the majority of reads represent redundant isoforms. **(C)** The number of unique internal exons, unique canonical introns and non-redundant isoforms resolved by single-molecule RNA CaptureSeq. Content from protein-coding genes (upper) and noncoding loci (lower) are shown separately and, for protein-coding genes, exons/introns belonging to noncoding isoforms or UTR variants (green) are distinguished from predicted ORFs (blue). (**D**) The proportion of captured bases on Hsa21 that are exonic, intronic or silent, according to Gencode (v19; left) RNA CaptureSeq (right).

At protein-coding loci on Hsa21, we identified 7,310 unique multi-exonic isoforms, of which 77% were novel, with respect to current annotations (a combined transcriptome catalog of Gencode v19, MiTranscriptome v2 and FANTOM5; **Figure 1C**; **Supplementary Table 5,6**). Novel isoforms encoded up to 2,365 possible ORFs that were not currently annotated, encompassing 291 novel coding exons and 845 novel ORF introns (**Figure 1C**; examples in **Figure S4**). Although these are predicted ORFs only, and the precise number retrieved is influenced by the choice of cutoff parameters (**STAR Methods**), this represents a considerable increase on existing annotations.

While this suggests that the protein-coding content of Hsa21 may be underestimated, most novel isoforms to protein-coding genes were noncoding isoform variants or possessed novel untranslated regions (UTRs). Alternative splicing of UTRs (both 5′ and 3′) was common and often highly complex (examples in **Figure S5**). Single-molecule sequencing was particularly useful for resolving UTR variants, since it does not suffer from sequencing 'edge effects' that impact short-read transcript assembly (Martin and Wang, 2011). In total, protein-coding loci on Hsa21 encoded 3,931 unique internal exons (34% novel) and 6,365 unique canonical introns (43% novel; **Figure 1C**; **Supplementary Table 5,6**).

At noncoding loci, we identified 1,589 isoforms across Hsa21, encompassing 1,663 unique internal exons (45% novel) and 3,210 unique canonical introns (55% novel; **Figure 1C**; **Supplementary Table 5,6**). Examples of rich isoform diversity at lncRNA loci were routinely resolved with targeted single-molecule sequencing (**Figure 1B**; **Figure S6**). In many instances, multiple partial lncRNA annotations were incorporated into single unified loci (**Figure S6A-C**), and the splicing of lncRNAs with neighboring protein-coding genes to form extensively spliced UTRs was also frequently observed (**Figure S6D**).

By extrapolating our Hsa21 annotations across the broader human genome, we can estimate the existence of 383,000 unique isoforms to protein-coding genes that encompass 147,000 possible ORFs. We predict noncoding gene loci to express 98,000 multi-exonic isoforms (2.1-fold increase by comparison to Gencode v26), incorporating 88,000 internal exons (1.2-fold) and 168,000 introns (1.7-fold). While we have profiled only three tissues, ensuring these are lower-bound estimates, it is clear that a large amount of transcriptional diversity remains unexplored.

### Universal alternative splicing of noncoding exons

To determine whether RNA CaptureSeq provided a complete profile of transcription on Hsa21, or whether further isoforms remain to be discovered, we generated discovery-saturation curves by incremental subsampling of short-read alignments (**STAR Methods**). The detection of protein-coding exons and introns approached saturation at a fraction of library depth, indicating that these were near-comprehensively sampled (**Figure 2A**; note that terminal exons were not considered in this analysis). While the detection of noncoding exons also approached saturation, the discovery of noncoding introns (and consequently additional noncoding isoforms) continued progressively toward maximum sequencing depth (**Figure 2A**). This indicates that, although the majority of exons were discovered, noncoding isoforms were not exhaustively resolved, even with the enhanced sensitivity afforded by RNA CaptureSeq.

**Figure 2.**
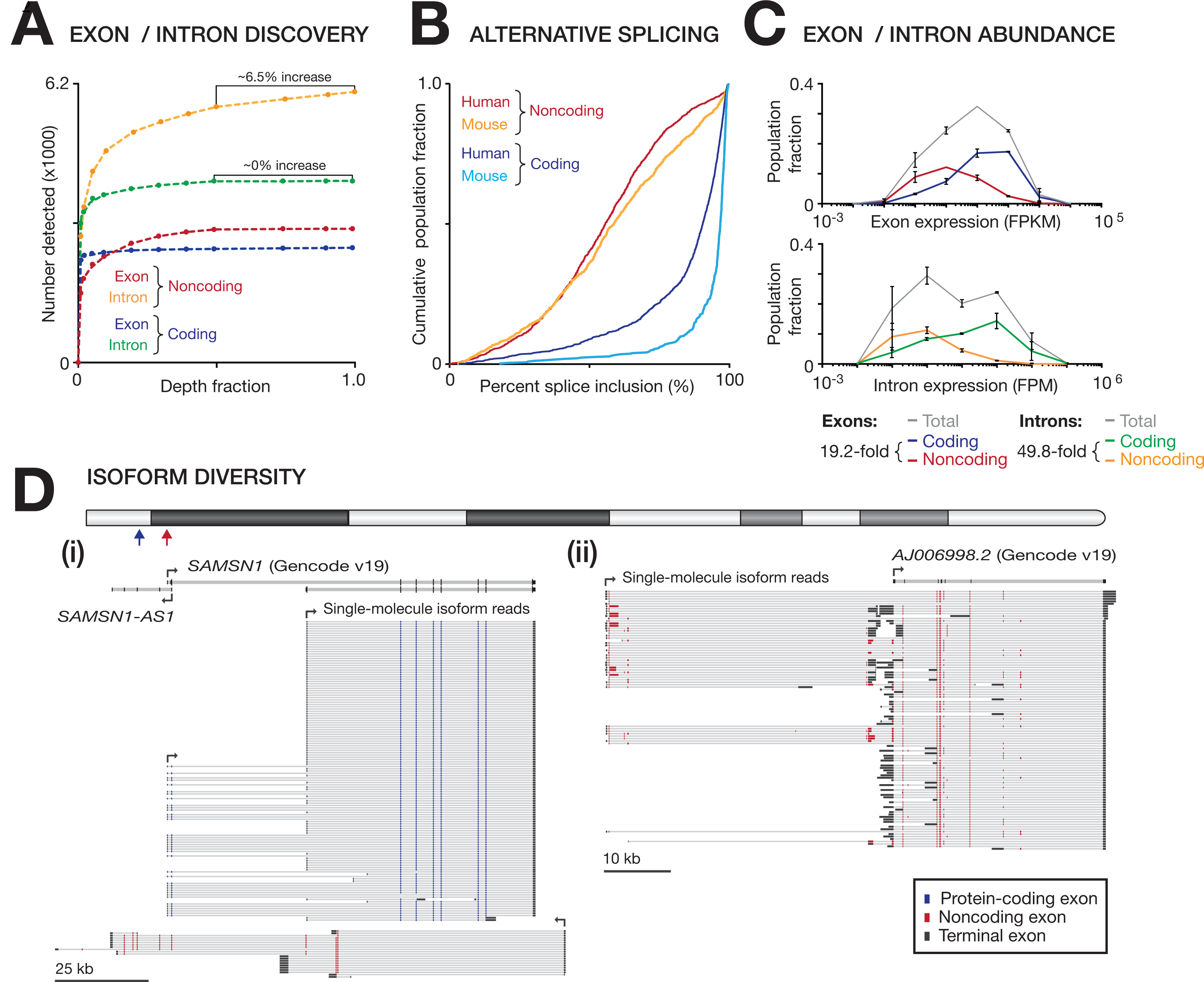
Universal alternative splicing of noncoding exons. (A) Discovery-saturation curves show the rate of detection for unique protein-coding/noncoding introns and unique internal exons (from the single-molecule Hsa21 transcriptome) relative to short-read sequencing depth. Depth fraction is relative to a combined pool of ~450 million short-read alignments to Hsa21 from testis, brain and kidney. **(B)** Cumulative frequency distributions show percent splice inclusion (PSI) scores for protein-coding/noncoding internal exons in human and mouse tissues. **(C)** Binned frequency distributions show abundances of protein-coding/noncoding internal exons and introns in human testis (mean + SD, *n* = 3). Grey line shows total exon/intron population. Median fold-difference between coding/noncoding populations is below. **(D)** Illustrative examples of isoforms resolved by single-molecule RNA CaptureSeq in human tissues. Annotated transcripts (Gencode v19) and mapped single-molecule isoform reads are shown at two loci: **(i)** the protein-coding gene *SAMSN1* and the noncoding antisense RNA *SAMSN1-AS1;* **(ii)** the lncRNA *AJ006998.2.* Internal exons are identified as protein-coding (blue) or noncoding (red), which includes untranslated exons at protein-coding loci. Terminal exons (black) were excluded from analyses.

Because each unique intron represents a different splicing event, this result suggests that alternative splicing generates a seemingly limitless diversity of noncoding isoform variants. To confirm this, we assessed the alternative splicing of internal exons according to Percent Splice Inclusion scores (PSI; **STAR Methods**). Unlike protein-coding exons (median PSI = 90.5%), almost all noncoding exons were alternatively spliced (median PSI = 55.5%; **Figure 2B; Figure S7**). The relative abundances of coding and noncoding introns, compared to exons, further supports this finding; we found that the relative difference between coding and noncoding introns (49.8-fold) is larger than for exons (19.2-fold; **Figure 2C**), reflecting the greater isoform diversity generated by enriched alternative splicing of noncoding RNAs. We use the phrase universal alternative splicing to mean that nearly every noncoding exon is subject to alternative splicing.

The distinction between coding and noncoding splicing is illustrated by comparison of the protein-coding gene *SAMSN1* to its antisense lncRNA (*SAMSN1-AS1*) and a nearby intergenic lncRNA (*AJ006998.2*). Whilst the majority of single-molecule *SAMSN1* transcripts correspond to one of just two mRNA isoforms for this gene, almost all *SAMSN1-AS1* and *AJ006998.2* transcript molecules represent unique isoforms (**Figure 2D**).

Universal alternative splicing was not limited to lncRNA exons but was similarly observed for untranslated exons at protein-coding loci (located in 5′ or 3′UTRs, or specific to noncoding isoforms; **Figure S8, S9**). For example, the 5'UTR of the protein-coding gene *CHODL* exhibited extensive alternative splicing (**Figure S10**). Splice acceptor sites at lncRNA and UTR exons harbored canonical splicing elements that were indistinguishable from those found at protein-coding exons (**Figure S8B**), confirming that noncoding exons are demarcated by *bona fide,* rather than cryptic splice sites.

Our analysis reveals a fundamental distinction between the organization of protein-coding and noncoding gene content. While the diversity of protein-coding isoforms is limited by the requirement to maintain an ORF, no such constraint is imposed on noncoding RNA, allowing the spliceosome to explore the full range of noncoding exon combinations to generate an effectively inexhaustible noncoding isoform diversity. It is worth noting here that this phenomenon is not limited to obscure noncoding RNAs encoded on Hsa21: using publicly available data (Cabili et al 2011; Clark et al 2015; Iyer 2015) we examined four well-known functional lncRNAs – *XIST*, *HOTAIR*, *GOMAFU* and *H19* – revealing rich isoform diversity and near-universal alternative splicing at each locus (**Figure S11**).

### Comparison of transcriptional landscapes in human and mouse

To establish whether the novel genes and isoforms unearthed on Hsa21 were conserved between human and mouse, and to determine whether noncoding exons are similarly enriched for alternative splicing, we performed short-read RNA CaptureSeq across mouse genome regions syntenic to Hsa21 (located on mouse chromosomes 10, 16 and 17; Waterson et al., 2002). We obtained transcriptional profiles of equivalent depth to human samples within matched mouse tissues (**Supplementary Table 7**), enabling comparison of the two transcriptomes at high-resolution (**STAR Methods**).

As for Hsa21, the syntenic regions of the mouse genome were pervasively transcribed, largely eroding intergenic desert regions (**Figure 3A**). Spliced transcripts encompassed 80.5% of targeted bases in mouse, compared to 88.4% on Hsa21 (**Figure 1D** vs **3B**). In mouse, 25.1% of targeted bases were retained as mature exons, compared to 16.7% in human, with the remaining fraction (55.4% vs 71.7%) removed as introns (**Figure 1D** vs **3B**). The larger fraction of bases represented in mature exons in mouse (1.5-fold) likely reflects compaction of the mouse genome via accelerated genetic loss (Waterson, et al., 2002; Vierstra et al., 2014; Yue et al., 2014), rather than higher gene content, with the mouse transcriptome assembly being somewhat smaller than human (77%, based on unique internal exon count; **Supplementary Table 8**).

**Figure 3.**
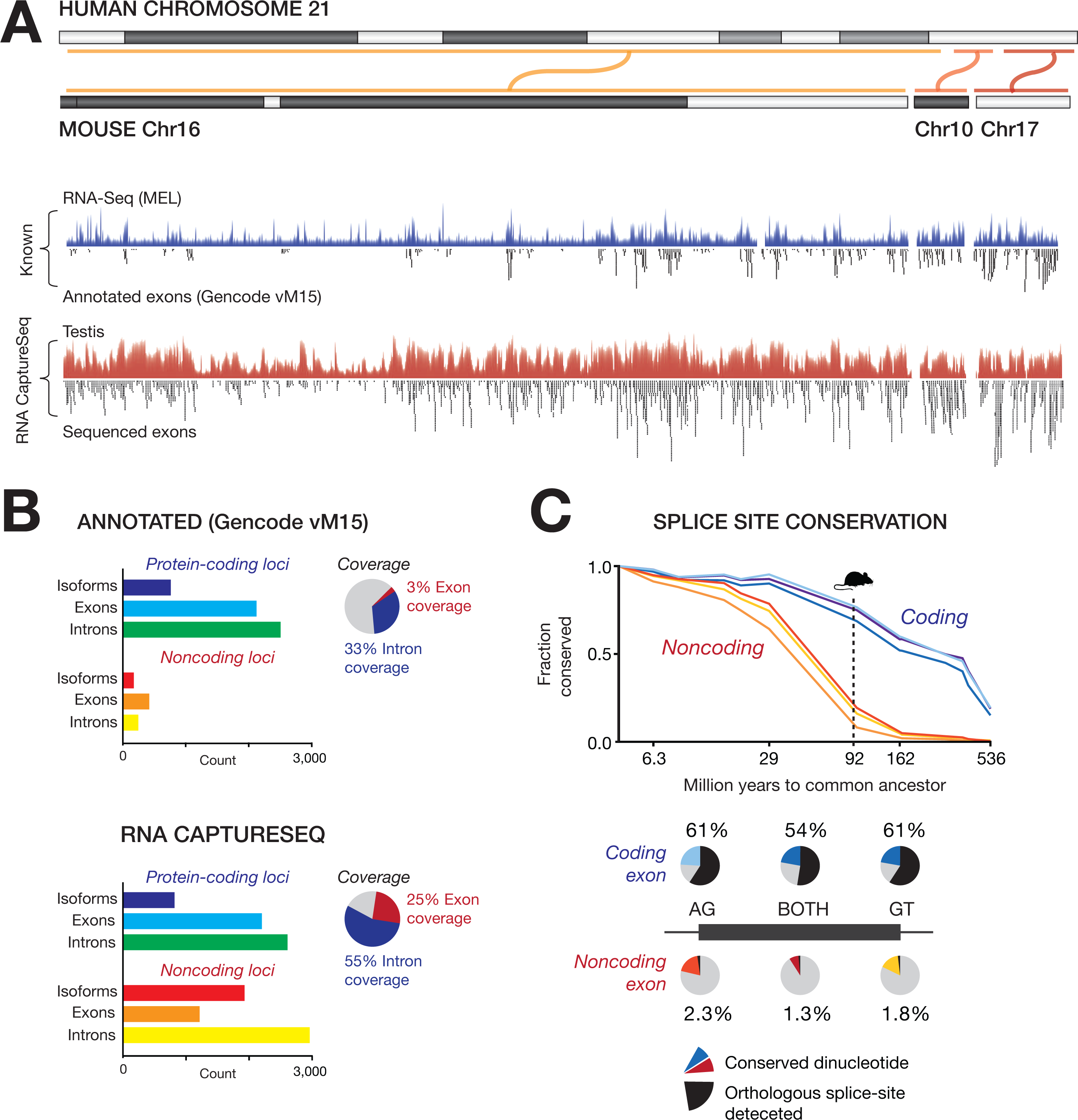
Transcriptional landscape of mouse syntenic regions and noncoding exon evolution. (A) Transcriptional activity recorded across regions of the mouse genome (Mmu16, Mmu17, Mmu10) syntenic to human chromosome 21 (Hsa21) by short-read RNA CaptureSeq. Normalized coverage (log scale) of short-read alignments is shown for testis (red), with traditional RNA-Seq (in MEL cells; navy) shown for comparison. Below coverage tracks are all unique internal exons (black) from transcripts in the Gencode (vM15) reference catalog and those assembled from RNA CaptureSeq data (testis, brain, kidney combined). **(B)** Bar charts show the number of protein-coding/noncoding unique internal exons, unique canonical introns and unique transcript isoforms in Gencode (vM15) annotations (upper) or assembled from RNA CaptureSeq (lower; brain, kidney and testis combined). Pie charts indicate the proportion of captured bases that are exonic, intronic or silent. (**C**; upper) The fraction of coding/noncoding splice-site dinucleotides (AG/GT/both) detected on Hsa21 that are conserved in other vertebrate genomes (arranged by million years to common ancestor). (**C**; lower) Pie-charts indicate the proportion of Hsa21 exons with splice-site dinucleotides that are conserved in the mouse genome (red/blue) and the proportion for which an equivalent splice site could also be detected in mouse RNA CaptureSeq libraries (black).

Although almost all protein-coding genes on Hsa21 have mouse orthologs, we observed a higher frequency of alternative splicing among human genes than their mouse counterparts: 69% of human protein-coding exons were classified as alternative (PSI < 95%) compared to just 31% in mouse (**Figure 2B**). As a result of this greater splicing diversify, human protein-coding genes had more internal exons (17%), introns (39%), isoforms (65%) than their corresponding mouse orthologs (**Supplementary Table 8**).

The *DYRK1A* gene, a leading candidate for autism and trisomy-21 phenotypes (Becker et al., 2014), provides an illustrative example of the increased splicing diversity distinguishing human genes from their mouse orthologs. While we found no novel exons or isoforms to the *Dyrk1A* gene in the mouse, we identified six novel internal exons in the human brain (in addition to all 13 currently annotated *DYRK1A* exons; **Figure 4A,B**). Extensive alternative splicing generated at least 11 novel *DYRK1A* isoforms, of which 10 comprise noncoding variants and one encodes a novel ORF with a N-terminal modification to the DYRK1A protein (**Figure 4A,B**; interestingly, an analogous N-terminal modification regulates subcellular localization of the DYRK4 paralog; Papadopoulos et al., 2011)).

**Figure 4.**
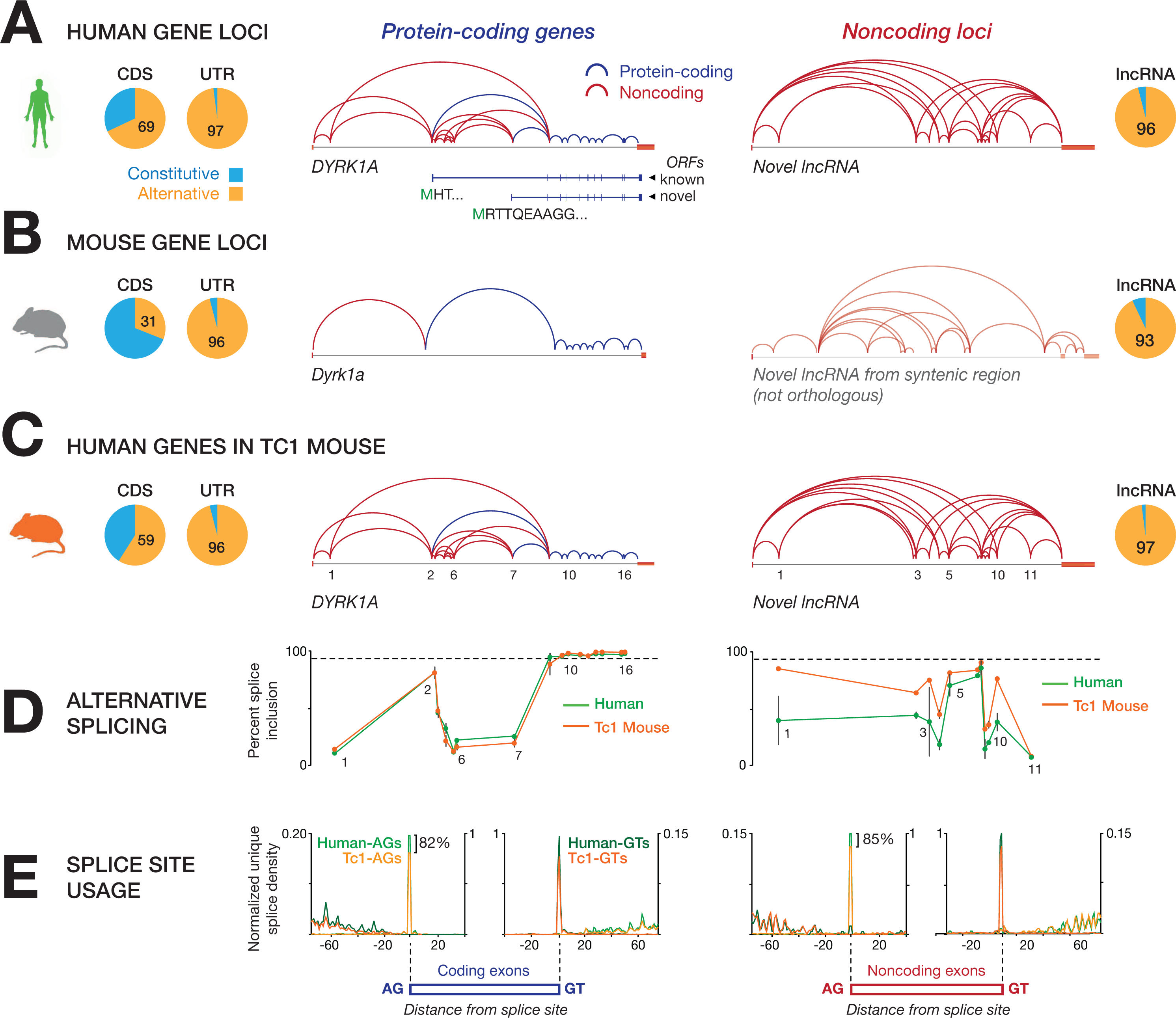
Splicing of human genes in the Tc1 mouse. **(A-C)** Exon-intron structures assembled for the protein-coding gene *DYRK1A* and a novel lncRNA locus from human-Hsa21 **(A),** mouse-syntenic regions **(B)** and on Hsa21 in the Tc1-mouse strain **(C).** Pie charts indicate the proportion of unique internal exons classified as constitutive (percentage splice inclusion; PSI > 95) or alternative, for protein-coding exons (CDS) and untranslated exons at coding loci (UTR) and lncRNA exons. **(D)** Plots show relative isoform abundance of *DYRK1A* and novel lncRNA loci in human and Tc1-mouse, as indicated by PSI values for each internal exon (aligned to exons in gene models above). PSI values shown for *DYRK1A* are measured from human brain libraries (mean ± SD, *n* = 2) and lncRNA from testis libraries (*n* = 3). **(E)** Density plots show global concordance of unique splice junction selection between human-Hsa21 (mean + SD, *n* = 2) and Tc1-Hsa21 libraries (*n* = 3) for human exons. Density values are normalized relative to human-Hsa21 libraries.

The majority of novel exons discovered in the mouse syntenic regions were noncoding (**Figure 3B**), and these were similarly subject to near-universal alternative splicing (**Figure 2B**). This indicates that the size and structure of the human and mouse transcriptomes are largely comparable, with each harboring large noncoding RNA populations that exhibit prolific alternative splicing.

Despite this similarity, we found that individual lncRNAs were largely divergent between the two lineages (**STAR Methods**). Nineteen percent of lncRNA splice-acceptor and 16% of splice-donor dinucleotides (AG/GT) were conserved between the human and mouse genomes. However, a corresponding splice site was found in mouse for fewer than 2% of human sites, implying that noncoding exon orthologs are rare (**Figure 3C**). Although they were poorly conserved relative to their protein-coding counterparts, noncoding exons exhibited internal sequence constraint (base on Vertebrate PhlyoP scores) comparable to annotated DNase hypersensitive or transcription factor binding sites, with flanking splice sites showing a further, though relatively modest, conservation enrichment (**Figure S12A,B**). These data indicate that, whilst similar in size and structure, human and mouse noncoding RNA populations are largely distinct, echoing the reported divergence of regulatory elements between mouse and human genomes (Vierstra et al., 2014; Villar et al., 2015).

### Human chromosome 21 expression and splicing in mouse cells

The Tc1 mouse strain is a model for trisomy-21 that carries a stable copy of Hsa21 (Yu et al., 2010). The Tc1 mouse has also been used to compare the human and mouse transcriptomes, enabling the regulatory contributions of human *cis*-elements and mouse *trans*-acting factors to be distinguished (Barbosa-Morais et al., 2012; Wilson et al., 2008). To investigate the regulation of transcriptome diversity, we performed shortread RNA CaptureSeq, targeting both Hsa21 and its syntenic mouse genome regions, in matched tissues of the Tc1 mouse (**Supplementary Table 9**; **STAR Methods**).

Most notably, the splicing profiles of genes encoded on Hsa21 were recapitulated in the Tc1 mouse as for human tissues, rather their mouse orthologs, where these were divergent. The *DYRK1A* gene again provides an illustrative example, with human-specific splice site selection and quantitative exon usage faithfully recapitulated on Hsa21 in Tc1 samples (**Figure 4A-D**). Globally, 87% of human-specific splice sites distinguishing human and mouse orthologs were also detected on Hsa21 in the Tc1 mouse (compared to 82% for shared splice sites; **Figure S13A**). Similarly, the alternative splicing frequency of human exons remained 2.1-fold higher than for mouse orthologs, and 88% of sites classified as alternative in human were also classified as alternative in Tc1 (compared to 39% in mouse; **Figure S13B,C**). When correlated according to the PSI profiles across all orthologous splice sites, we found that human, mouse and Tc1 samples clustered according to chromosomal, rather than organismal, origin (**Figure S14C**). Together these data confirm the central importance of local *cis* sequence elements in defining exon boundaries and alternative exon inclusion.

The structure and splicing of lncRNAs encoded on Hsa21 was also precisely recapitulated in the Tc1 mouse, despite the absence of mouse orthologs. The majority (85%; **Figure 4E**) of human noncoding splice sites were also detected in Tc1 and noncoding exons were again near-universally alternatively spliced (98%; **Figure 4A-C**). Furthermore, quantitative splice site usage and relative noncoding isoform abundance was maintained as for human tissues (**Figure 4D**), indicating that the local Hsa21 sequence is sufficient to establish splice site position and regulate the proportional inclusion of noncoding exons by alternative splicing.

In contrast to splicing, we observed a global deregulation of expression in the Tc1 mouse, as assessed by principle component analysis or rank-correlation clustering (**Figure S14A,B**). This effect is best illustrated at intergenic regions flanking the *NCAM2* locus, where numerous lncRNA genes that are silenced in the human brain become deregulated, resulting in aberrant expression in the Tc1 mouse brain (**Figure S15A**). In fact, the expression of human protein-coding genes encoded on Hsa21 was more similar to expression of their mouse orthologs in corresponding mouse tissues than to the expression of the same human genes encoded on Hsa21 in the Tc1 mouse (**Figure S14A,B**). This deregulation was restricted to the human chromosome, with tissue-specific expression profiles still maintained across syntenic regions of the mouse genome in Tc1 mouse (**Figure S15B**).

This analysis appears to highlight a distinction in the evolution of expression and splicing regulation. The deregulation of human gene expression in mouse cells is consistent with the reported divergence of human and mouse regulatory elements, including enhancers and transcription factor binding sites (Vierstra et al., 2014; Villar et al., 2015). In contrast, splicing profiles were largely recapitulated on Hsa21 in the Tc1 mouse, as has been observed previously for protein-coding exons (Barbosa-Morais et al., 2012). This implies that splicing is largely regulated by *cis* elements in the local chromosome sequence (Barash et al., 2010) and can be correctly interpreted by the mouse spliceosome. Our findings imply that the splicing code is so highly conserved that human-specific exons and noncoding RNAs without orthologs in mouse are correctly spliced. This demonstrates deep conservation of the lexicon that governs splicing, even whilst the isoforms produced undergo rapid diversification and turnover.

## DISCUSSION

To overcome the expression-dependent bias in RNA-Seq, which impedes discovery and characterization of low abundance transcripts (Hardwick et al., 2016), we performed targeted RNA sequencing across human chromosome 21. The combination of single-molecule and short-read RNA CaptureSeq enabled accurate resolution of isoform diversity and quantitative analysis of alternative splicing at unprecedented depth.

Noncoding loci are, contrary to the impression from more shallow surveys (Cabili et al., 2011; Derrien et al., 2012), enriched for alternative splicing, with noncoding exons being near-universally classified as alternative. This finding is consistent with previous reports of lower splicing efficiency and U2AF65 occupancy among lncRNAs than mRNAs, features independently correlated with heightened alternative over constitutive splicing (Mele et al., 2017, Mukherjee et al., 2017, Tilgner et al., 2012).

Therefore, while protein-coding genes are constrained by the requirement to maintain an ORF, it appears that no similar constraint is imposed on noncoding RNA. This suggests that noncoding exons are functionally modular, operating as discrete cassettes that are recombined with maximum flexibility. One can envision a scenario where individual noncoding exons interact independently with other biomolecules (proteins, RNAs and/or DNA-motifs), organizing these around the scaffold of a noncoding transcript. In this way, alternative isoforms could assemble different collections of binding partners to dynamically regulate cellular processes. The distinction between protein-coding and noncoding RNA was also evident when comparing exons within ORFs to untranslated regions of coding loci (located in 5′ or 3′UTRs, or specific to noncoding isoforms), implying similar modularity in the functional architecture of untranslated regions.

Low expression is often cited as evidence against the functional relevance of novel transcripts, such as the wide variety of rare noncoding isoforms identified in our survey. However, while weakly expressed genes/isoforms are unlikely to fulfill structural or metabolic functions, there are precedents for these fulfilling regulatory roles. For example, Marinov et al. (2014) found the mRNAs of expressed transcription factors to be present, on average, at just ~3 copies per cell within a homogenous cell line (GM12878). Single-cell studies routinely reveal rare cell types within human tissues on the basis of their unique gene expression profiles. These may be represented by just a few cells within a community of hundreds or thousands of cells (Grun et al., 2015; La Manno et al., 2016; Muraro et al., 2016). Given that mRNAs encoding regulatory molecules like transcription factors are expressed at only a few copies per cell, the regulatory factors that define the identity of rare cell types are expected to be present at very low frequencies in whole-tissue transcriptomes. It would be premature, therefore, to discount the relevance of any transcript on the basis of low abundance alone.

Moreover, while not every rare isoform is necessarily important, and the promiscuous splicing we observed might simply reflect a lack of selective pressure, noncoding RNAs collectively form a large reservoir of transcriptional diversity from which molecular innovations might evolve and new genes may be born (Kaessmann, 2010; Toll-Riera et al., 2009; Wu et al., 2011; Xie et al., 2012). The use of alternative splicing to generate noncoding transcriptional diversity and, thereby, drive gene evolution is consistent with the divergence of noncoding exons reported here. By contrast, the splicing code that governs such transcriptional diversity remains closely conserved between human and mouse.

Despite concerted efforts over the past decade, we are yet to achieve a complete census of human gene expression. Even our use of targeted single-molecule RNA sequencing was insufficient to resolve the full complement of noncoding isoforms encoded on Hsa21. Instead, we found a seemingly limitless diversity of noncoding isoforms. Given the range of combinatorial possibilities, we suggest that the noncoding RNA population may be inherently plastic, and that there does not exist a finite list of noncoding isoforms that can be feasibly catalogued.

## AUTHOR CONTRIBUTIONS

T.R.M. and M.E.B. conceived the project and designed experiments, with advice from J.S.M., L.K.N and M.E.D. M.E.B., M.B.C., J.C. and J.B. performed capture enrichment and library preparations for short-read sequencing. M.E.B. and J.B. performed PCR validations or assembled transcripts. J.B. and T.H. performed long-read sequencing, overseen by T.A.C. I.W.D. and E.T. performed bioinformatic analyses. I.W.D. and T.R.M. prepared the manuscript with support from M.E.B., J.B., M.B.C., L.K.N. and J.S.M. T.R.M., M.E.D., L.K.N. and J.S.M. provided funding. Correspondence should be addressed to T.R.M.

## ACKNOWLEDGEMENTS

The authors acknowledge the following funding sources: an Australian National Health and Medical Research Council (NHMRC) Project Grant (APP1062106 to T.R.M.), NHMRC Australia Fellowship (631668 to J.S.M.), an NHMRC Early Career Fellowship (APP1072662 to M.B.C.), an EMBO Long Term Fellowship (ALTF 864-2013 to M.B.C.), the the Australian Research Council (Special Research Initiative in Stem Cell Science to L.K.N.) and the generous support of the Paramor family (to T.R.M.). The contents of the published material are solely the responsibility of the administering institution, a participating institution or individual authors and do not reflect the views of NHMRC or ARC. The authors thank the ENCODE consortium for the provision of data; data were employed in accordance with the data-release policy.

## STAR METHODS

### CONTACT FOR REAGENT AND RESOURCE SHARING

Further information and requests for resources and reagents should be directed to and will be fulfilled by the Lead Contact, Tim Mercer.

### EXPERIMENTAL MODEL AND SUBJECT DETAILS

#### Human samples

Total RNA samples from healthy adult tissues were acquired from a commercial vendor (Ambion Human RNA Survey Panel). Ambion certifies that all of human-derived materials have been prepared from tissue obtained with consent from a fully informed donor or a member of the donor’s family. Two replicate samples for human brain and kidney, and three replicates for testis were analyzed.

#### Mouse samples

All animals were handled, housed, and used in the experiments in accordance with protocols approved by the Animal Ethics Committee of The University of Queensland, Brisbane, Australia. Three male mice (C57BL/6J x 129S8/SvEv) and 3 Tc1 female breeders were imported from the Jackson Laboratory, Maine, USA. 10 x F1 pups produced from paired mating were toed for identification and genotyping. Genotyping was performed on gDNA extracted from toe tissues using primers specific to human and mouse *JAM2* genes. Three Tc1 F1 males and 3 WT littermates were sacrificed between 6 and 9 weeks of age, and testis, kidney and brain tissues were immediately harvested in TRIzol reagent (Ambion), snap-frozen on dry ice and stored at −80°C until processing.

#### K562 cells

The human lymphoblast cell line, K562, was obtained from the American Type Culture Centre (ATCC). Cells were not independently verified or tested for mycoplasma. Cells were cultured according to Coriell Institute’s growth protocols and standards. Briefly, K562 cells were cultured in RPMI 1640 medium (Gibco) supplemented with 10% FBS at 37 °C under 5% CO_2_. Cells (passage 9 or 10) were grown to ~80% confluence before RNA extraction.

### METHOD DETAILS

#### Total RNA extraction

##### Mouse samples

Tissue samples (brain, kidney and testis), harvested in trizol and snap-frozen (see above), were defrosted on ice and 125 mg MicroBeads (MoBio) were added to sample tubes. Tissues and cells were disrupted by 2 cycles of 45 sec at 6500 rpm on a tissue homogenizer (Precellys) connected to a cooling system (Cryolis) keeping the samples temperature < 4°C during the cell lysis procedure. Bead-free cell lysates were transferred to a new 1.5 mL tube and RNA was immediately harvested, as above. Brain RNA was extracted a second time in TRIzol due to high lipid contamination present after the first extraction.

##### K562 cells

Total RNA was extracted in 3mL TRIzol reagent (Ambion). Extraction of the aqueous phase was performed using chlorophorm and RNA precipitation was achieved with isopropanol. RNA pellets were washed in ice-cold 75% EtOH and allowed to dry before suspension in DEPC-treated H_2_O.

#### Capture enrichment

Oligonucleotide probes targeting the entire non-repetitive portion of Hsa21 (hg19) and its syntenic regions in the mouse (mm10) genome, obtained using the UCSC Genome Browser liftOver tool, were designed and synthesized by Roche/NimbleGen. In total, the array targeted 25.3 Mb encompassing 0.87% of the human genome and 20.3 Mb encompassing 0.77% of the mouse genome.

Total RNA samples were assessed for potential gDNA contamination by PCR and accordingly treated with Turbo DNAse (Ambion). Samples were spiked with ERCC RNA spike-in controls (Jiang et al., 2011) to a final 1% concentration and rRNA-depleted using the Ribo-Zero rRNA removal magnetic kit (Epicentre). cDNA libraries were prepared from rRNA-depleted samples using the Illumina TruSeq stranded mRNA low-template kit. Pre-capture LMPCR amplified libraries were purified using AMPure XP beads (Beckman Coulter Genomics), and successful library preparation was validated using a DNA 1000 kit (Agilent) on a Bioanalyser. Average library sizes were ~280-310 bp.

Capture hybridization was performed in 96-well plates, with hybridization probes incubated at 47 °C for 64-77 h. Post-capture LMPCR amplification was performed for 17 cycles. Amplified post-capture libraries were purified using AMPure XP beads, and validated using a DNA 1000 kit on a Bioanalyzer. Samples that retained unincorporated primers were cleaned a second time using a QIAquick PCR purification kit (Qiagen), and successful primer removal was validated on a Bioanalyzer. For a detailed protocol see (Mercer et al., 2014).

#### PacBio SMRTbell library preparation and single-molecule sequencing

A total of 4.5 μg of captured full-length cDNA was subjected to size fractionation using the Sage Science BluePippin system into four size bins (0-2kb, 1-3kb, 3-6kb and 4-10kb). Eluted size fractions were subsequently re-amplified and purified with AMPure PB beads. Size distributions of the fractions were checked for quality on a 2100 BioAnalyzer (Agilent).

Approximately 10 μg of purified amplicon was taken into Iso-Seq SMRTBell library preparation (https://pacbio.secure.force.com/SamplePrep). Separate SMRTbell libraries were generated for each of the four size bins (**Supplementary Table 10**). Two of the SMRTBell libraries (3-6kb and 4-10kb) were size-selected again using the Sage Science BluePippen system to remove trace amounts of small inserts. A total of 24 SMRT Cells (6 cells for each of the size-selected SMRTBell library) were sequenced on the PacBio *RS II* platform using P6-C4 chemistry with 3 to 4 hour acquisition time.

#### Short-read (Illumina) sequencing

Short-read sequencing libraries were prepared with the Illumina TruSeq stranded mRNA low-template kit. Libraries were sequenced at the Garvan Institute (Sydney, NSW, Australia) on the Illumina HiSeq 2000 or Illumina HiSeq 2500 platforms, generating 2 × 101bp and 2 × 125bp paired-end sequencing, respectively.

### QUANTIFICATION AND STATISTICAL ANALYSIS

#### Validation of Hsa21 RNA CaptureSeq in K562 cells

To establish the Hsa21 CaptureSeq approach we first carried out short-read sequencing (Illumina HiSeq 2000) on Hsa21-enriched libraries from the human K562 cell-type (*n* = 3) and compared these to K562 samples analyzed in parallel by conventional RNA-Seq (*n* = 3; **Supplementary Table 1**).

Sequencing libraries were trimmed using TrimGalore (https://www.bioinformatics.babraham.ac.uk-/index.html; default parameters), then aligned to a combined index of the hg19 reference genome and ERCC spike-in sequences. Alignment was performed by STAR (Dobin et al., 2013) with the following custom parameters:

--twopassMode Basic --outFilterIntronMotifs RemoveNoncanonical --alignIntronMin 20 --alignIntronMax 500000 \

On-target alignment rates were calculated by determining the number of uniquely mapped reads (MapQ=255) within Hsa21, as a fraction of all uniquely mapped reads across hg19. The average fold-increase in on-target rate for CaptureSeq samples, relative to non-captured samples, provides an estimate of the coverage enrichment achieved by capture (**Supplementary Table 1**).

Relative abundances (in FPKM) of ERCC RNA spike-ins were determined using RSEM (Li and Dewey, 2011), allowing the accuracy of transcript quantification to be assessed in captured/non-captured samples (**Figure S2A,B**). Progressively decreasing enrichment at very high ERCC concentrations was observed due to saturation of CaptureSeq oligonucleotide baits, but is unlikely to affect transcripts within the physiological range of gene expression (Clark et al., 2015; Mercer et al 2014).

#### Single-molecule Hsa21 transcriptome survey

To avoid potential artifacts associated with *de novo* transcript assembly, we analyzed Hsa21-enriched cDNA from human testis by deep single-molecule PacBio sequencing. Testis was selected because it provides a broad survey of gene-content encoded on Hsa21 (Soumillon et al., 2013).

The PacBio Iso-Seq “classify” protocol was used to generate full-length, non-chimeric (FLNC) reads (https://github.com/PacificBiosciences/cDNA_primer). Briefly, for each sequencing ZMW, a circular consensus sequence (CCS) was generated. Each CCS sequence was identified as full-length, non-chimeric if (1) both the 5′ and 3′ cDNA primer and the polyA tail preceding the 3′ primer was identified at the two ends; and (2) the sequence does not contain cDNA primers or polyA tails in the middle of the sequence (indication of library artifacts). Because FLNC reads are supported by the presence of a polyA tail, they are considered to represent mature transcript isoforms. Resolution of both sequencing primers ensures that FLNC reads have been sequenced in completeness. However, the possibility of degradation due to the use of template switching without 5′ cap capture means that some FLNC reads may not be biologically full-length.

FLNC reads were mapped to the human reference genome (hg19) using GMAP (Wu and Watanabe, 2005) and reads that mapped to Hsa21 with maximum mapping confidence (MapQ=40) were retained. In total, 387,029 reads aligned to Hsa21, constituting 910 Mb of usable transcript sequence concentrated in ~1.5% of the genome (**Figure S3A,B**; **Supplementary Table 3**; **Supplementary Table 10**).

To retain only high-quality multi-exonic single-molecule transcripts, we next discarded: (1) transcripts with < 3 exons; (2) transcripts that contained one or more non-canonical introns (based on AG/GT splice sites); (3) any transcript that was entirely contained as a partial fragment of a longer transcript or an annotated transcript (based on internal exon-intron-exon chain); (4) any transcript with the following CuffCompare classifier: e, i, o, p, r, s (Trapnell et al., 2012). After filtering, we retrieved 101,478 full-length multi-exonic transcripts (**Figure S3B; Supplementary Table 3**). To our knowledge, this represents the deepest singlemolecule transcriptome survey of a large genome region ever performed.

Transcripts were next assessed for overlap with known protein-coding genes. Transcripts that shared exonic overlap (on the same strand) with a known gene but not an identical internal exon-intron-exon chain were considered to represent novel isoforms of known genes. Redundant transcripts (sharing identical internal exon-intron-exon chains) were collapsed.

These were classified as protein-coding or noncoding via the Coding Potential Assessment Tool (Wang et al., 2013; longest ORF ≥ 100 codons, coding potential score ≥ 0.99). For coding transcripts we used TransDecoder to predict ORFs. Redundant ORFs and fragments fully contained within longer ORFs were discarded. ORFs were overlapped with full-length transcripts to distinguish coding exons from noncoding exons belonging to coding loci, including spliced UTR exons and exons incorporated exclusively into noncoding isoform variants at coding loci.

Transcripts that shared no exonic overlap with a known protein-coding gene (Gencode v19) and lacking a predicted ORF (CPAT; longest ORF < 100 codons, coding potential score < 0.99) were classified as lncRNA transcripts. Redundant transcripts (sharing identical internal exon-intron-exon chains) were collapsed.

#### Short-read Hsa21 RNA CaptureSeq in human tissues

To obtain quantitative information about transcripts in our single-molecule Hsa21 profile, short-read sequencing (Illumina HiSeq 2500) was performed on duplicate Hsa21-enriched libraries from brain, kidney and testis. Sequencing libraries were trimmed and aligned as described above (see above). On-target alignments rates were calculated as above (see **3.1**) and reads that were uniquely mapped (MapQ=255) within Hsa21 were retained for further analysis. At least 100 million alignments to Hsa21 were obtained for each tissue (**Supplementary Table 4**).

Because *de novo* transcript assembly from short sequencing reads may generate incorrect transcript models, especially for low-abundance isoforms (Hardwick et al., 2016), we did not perform transcript assembly. For all analyses of human tissues, the single-molecule PacBio transcriptome described above (see above) was used as a reference, with spliced-short read alignments used to quantitatively evaluate expression and splicing of PacBio transcripts.

The majority of spliced short-read alignment junctions were concordantly mapped to an intron in our single-molecule Hsa21 transcriptome profile (**Figure S3B**). Likewise, the majority of unique internal exons in our single-molecule Hsa21 transcriptome were correctly detected (both boundaries specified by ≥ 3 spliced short-read alignment termini) in at least one tissue (**Figure S3C**). This concordance with more accurate short-reads (Conesa et al., 2016) suggests that single-molecule isoforms were, in general, correctly aligned to Hsa21, and their internal exon-intron-exon architecture was correctly resolved.

#### Discovery saturation curves

To generate discovery-saturation curves, short-read alignments from brain, kidney and testis were combined, then incrementally subsampled from a maximum of ~450 million alignments. We assessed the detection of introns and exons within our single-molecule Hsa21 transcriptome by short-read alignments at each depth-increment. The analysis was limited to unique canonical introns and unique internal exons (as opposed to terminal exons) since these should be precisely demarcated by short-read alignment junctions at either end. An internal exon from our transcriptome was considered to be detected within a given library if both boundaries were specified by ≥ 3 spliced short-reads. An intron from our transcriptome was considered to be detected within a given library if it was spanned by ≥ 3 spliced short-read alignment junctions, mapping exactly to its 5′ and 3′ ends.

#### Percent Splice Inclusion (PSI) Values

To further assess alternative splicing, we calculated a Percent Splice Inclusion (PSI) score for each internal exon in our Hsa21 transcriptome, as has been done previously (Barbosa-Morais et al., 2012; Tilgner et al., 2015). For each exon, we counted spliced reads that used its 5′ splice site (**In**_1_) and 3′ splice site (**In**_2_) and spliced reads that spanned/skipped the exon (**Out**). The PSI value for that exon was then calculated as:

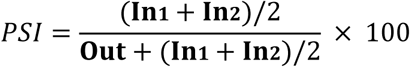

We confirmed that the occurrence of lower inclusion frequencies among noncoding exons (median PSI = 55.5%) than protein-coding exons (median PSI = 90.5%; **Figure 2B**) was not simply caused their lower gene expression, since exon-PSI and overall gene expression metrics were independent from one and other (**Figure S7**). Applying a threshold of PSI > 95% to distinguish constitutive from alternatively spliced exons, as has been done previously (Barbosa-Morais et al., 2012; Tilgner et al., 2015), almost all noncoding exons (97%) would be classified as alternative. We note that the use of a single, essentially arbitrary, PSI threshold is simplistic. However, this difference was indicative of a clear, population-wide depletion of PSI scores among noncoding exons, relative to their protein-coding counterparts, with a clear difference observable at any chosen threshold (**Figure 2B**).

The use of PacBio reads as a scaffold for calculating PSI scores ensures we are working with accurate transcripts (free of potential artifacts from transcript assembly). However, PacBio reads are less suitable for quantitative analyses because (i) size fractionation and heavy PCR amplification during the PacBio library preparation may distort transcript abundances within the population and (ii) saturating read-depth is required for accurate analyses of splicing. Therefore we consider our combined approach to be the most reliable way to assess alternative splicing. PSI scores were also calculated independently using only spliced PacBio long-read alignments, rather than spliced short-reads. This analysis produced similar results (**Figure S9**).

#### Comparison of human and mouse transcriptomes

To perform a fair comparison of transcriptome dimensions between human and mouse profiles, we generated *de novo* transcriptome assemblies for each, at matched depth, rather than using the single molecule human transcriptome (which would skew the comparison, since long-read sequencing was not performed on mouse tissues). While we note that *de novo* assembly may produce some spurious transcript models, the comparison of transcriptome dimensions between human and mouse is fair, since biases should affect both assemblies equally. Additionally, because assemblies were generated without the support of reference gene catalogs, they are not affected by discrepancies in the quality/completeness of human/mouse genome annotation.

After aligning reads to hg19 or mm10 with STAR and retaining only on-target, uniquely mapped reads (see above), libraries were subsampled to achieve a maximum matched depth of ~100 million alignments per tissue (**Supplementary Tables 4,7**). Assemblies were generated from these matched libraries using StringTie (Pertea et al., 2015). Assemblies from replicates for each tissue were merged using Cuffmerge (Trapnell et al., 2012). Full-length assembled transcript sequences were filtered and classified as above (see above).

#### Conservation analyses

To assess genomic-level sequence constraint we calculated average per-base PhyloP scores from 100 Vertebrate MultiZ alignments, obtained via the UCSC Genome Browser, within noncoding exons. Intergenic and intronic sequences of equivalent size were selected at random and used to provide a measure of neutral/background conservation. In addition to noncoding exons, we examined noncoding splice site dinucleotides (AG/GT), which were more strongly conserved than internal exon sequences, as well as annotated transcription factor binding sites and DHS sites on Hsa21 (**Figure S12**).

While PhyloP scores provide a useful measure of constraint on genomic sequence (inferred by comparison to other vertebrate genomes), our targeted transcriptome survey of syntenic human-mouse genome regions, allowed us to evaluate whether conserved exonic splice site nucleotides are actually transcribed and/or used as splice sites in both lineages. To do so, we lifted splice-site dinucleotides (AG/GT) demarcating human noncoding exons to the mouse genome using the UCSC liftOver tool. If a human splice-site dinucleotide was found to be conserved in the mouse genome, we tested to see if an equivalent junction could be found among mouse CaptureSeq alignments. While ~15% of human noncoding splice site nucleotides are conserved in mouse at the genomic level, a corresponding splice site was found in mouse for fewer than 2% of human sites.

#### Analysis of Hsa21 splicing and expression in the Tc1 mouse

The Tc1 mouse strain is a model for trisomy-21 that carries a stable copy of Hsa21 (Yu et al., 2010). We applied short-read RNA CaptureSeq (Illumina HiSeq 2500) to provide saturating coverage of both the Hsa21 transcriptome and syntenic mouse chromosome regions in Tc1 mouse tissues (brain, kidney and testis; **Supplementary Table 9**).

To assess alternative splicing we calculated PSI scores (see above) for protein-coding and noncoding exons expressed on Hsa21 or the mouse syntenic regions in Tc1 tissues. We compared PSI values at Tc1 exons to their equivalent exons detected in human or mouse, and between orthologous exons from the two species (**Figure S13**). To perform global comparisons of alternative splicing profiles among orthologous human/mouse exons, we clustered human/mouse/Tc1 mouse tissues according to their PSI scores using either principle component analysis or rank-correlation clustering (**Figure S14**). We did the same for gene expression, using FPKM values for orthologous genes rather than PSI scores.

### DATA AND SOFTWARE AVAILABILITY

Raw sequencing data and a combined transcriptome annotation are available via the NCBI Gene Expression Omnibus (GEO) under the following accession: GSE99637.

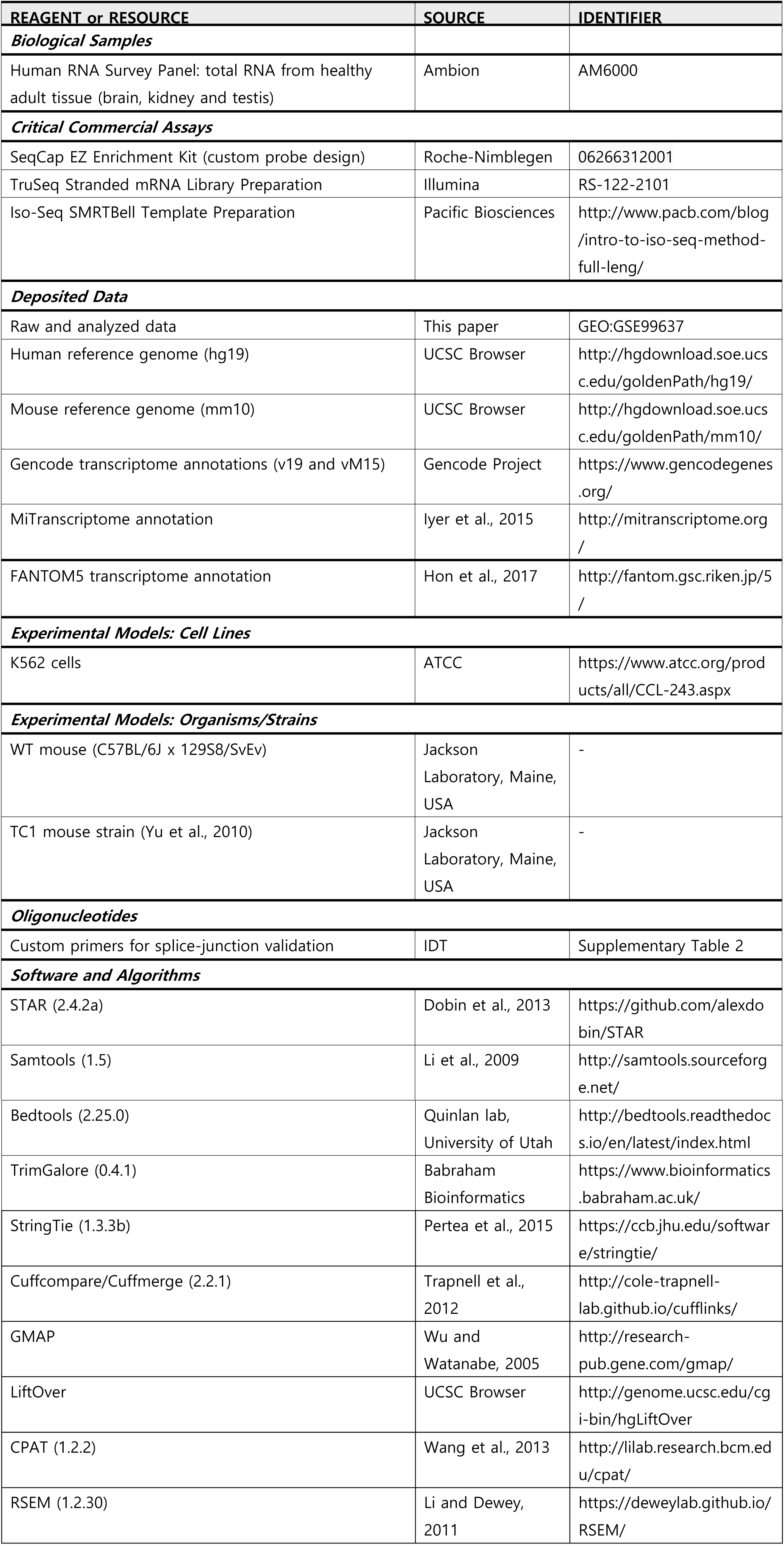
KEY RESOURCES TABLE

## SUPPLEMENTARY MATERIALS

**Figure S1. Related to Figure 1.**
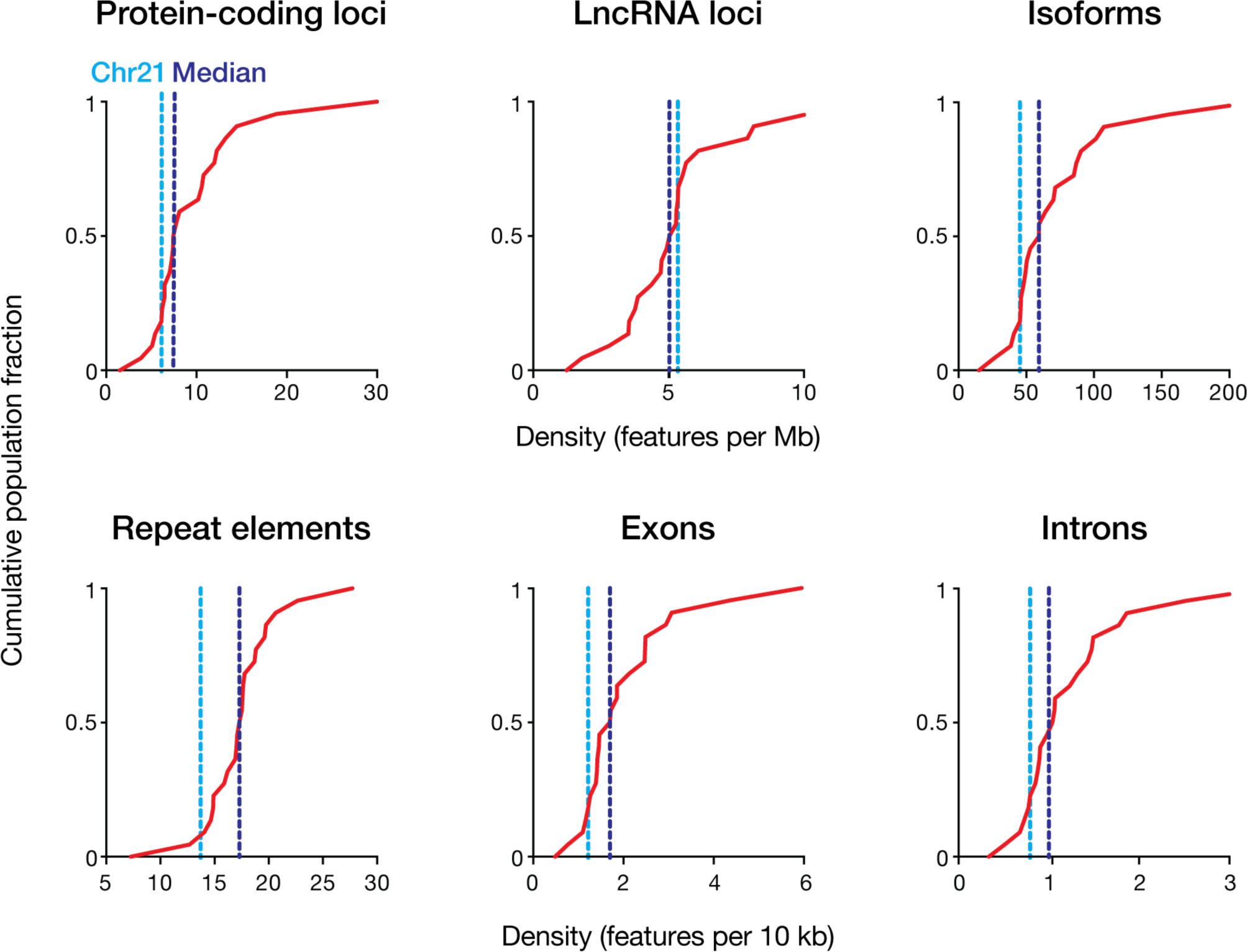
The distribution of genome features on Hsa21 is typical of the broader human genome. In order to reach the lower limits of the transcriptome, we surveyed gene expression from a single human chromosome with RNA CaptureSeq. Hsa21 was selected because it is the smallest human chromosome (48 Mb; ~1.5% of the genome). To confirm that Hsa21 is typical of the broader genome, we determined the density of protein-coding genes (Gencode v19), lncRNA genes (Gencode v19), repeat elements (RepeatMasker), transcript isoforms, exons and introns (Gencode v19) on Hsa21 and the other human autosomes. Cumulative frequency distribution plots indicate the density of each feature (features per Mb) across all human autosomes. Dashed lines indicate the density of features encoded on Hsa21 (light blue) and median feature density for all autosomes (dark blue).

**Figure S2. Related to Figure 1.**
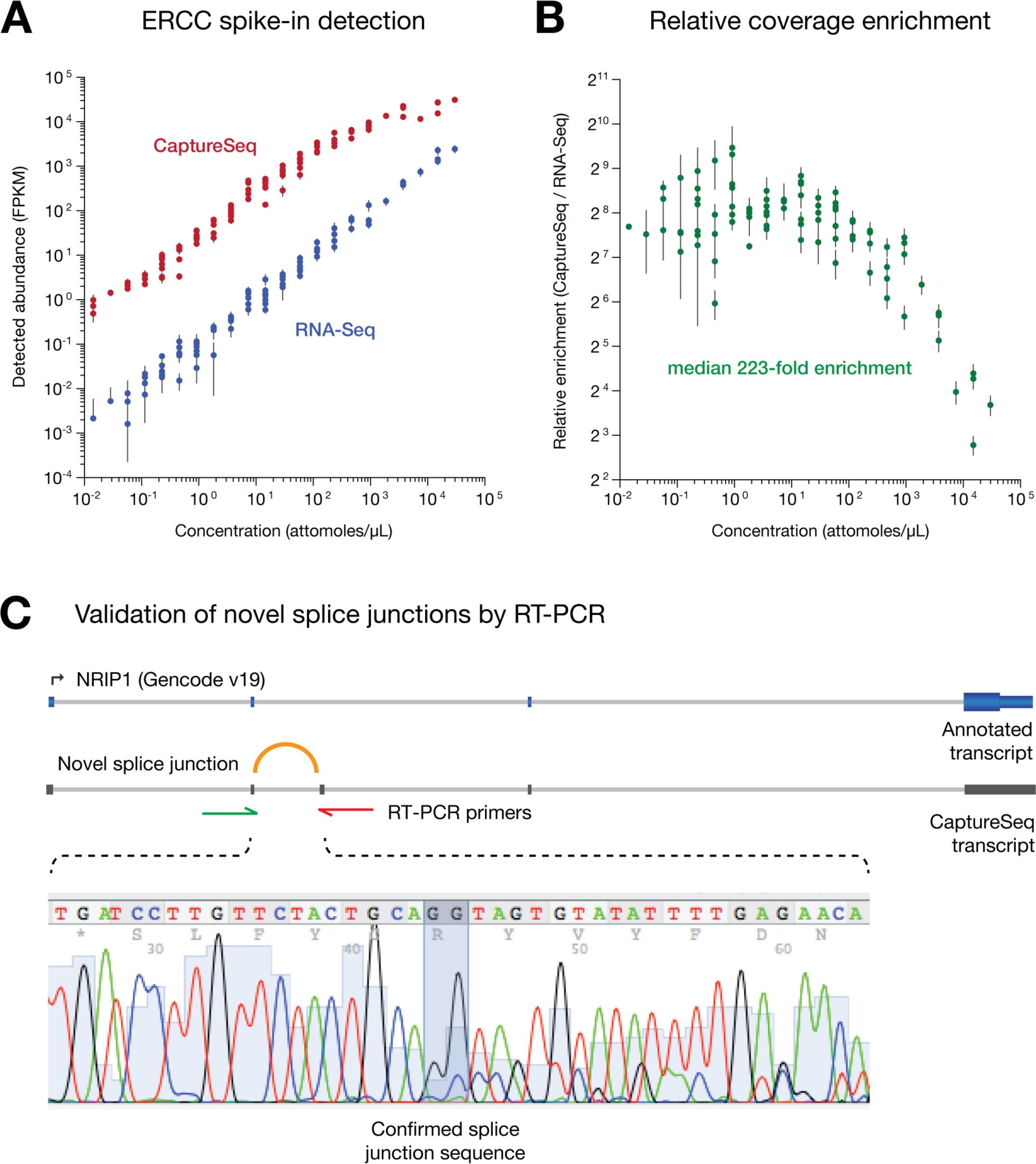
Performance of Hsa21 RNA CaptureSeq in human K562 cells. To validate the Hsa21 CaptureSeq method, we first performed short-read sequencing on Hsa21-enriched cDNA libraries from the human K562 cell-type and compared these to K562 samples analyzed in parallel by conventional RNA-Seq. **(A)** Abundances of ERCC RNA spike-in controls, relative to their input concentrations, as detected by conventional short-read RNA-Seq and RNA CaptureSeq in K562 samples (mean ± SD, *n* = 3). **(B)** Relative fold-enrichment achieved by RNA CaptureSeq for each ERCC standard. Variable enrichments observed at low input concentrations is the result of sporadic sampling when coverage is low. Progressively decreasing enrichments at high input concentrations is due to saturation of CaptureSeq probes. **(C)** The legitimacy of 16/20 (randomly selected) novel splice-junctions identified by CaptureSe were confirmed by RT-PCR and Sanger sequencing (**Supplementary Table 2**). To illustrate this approach, a single example is shown, in which a novel splice junction in the gene *NRIP1* was verified.

**Figure S3. Related to Figure 1.**
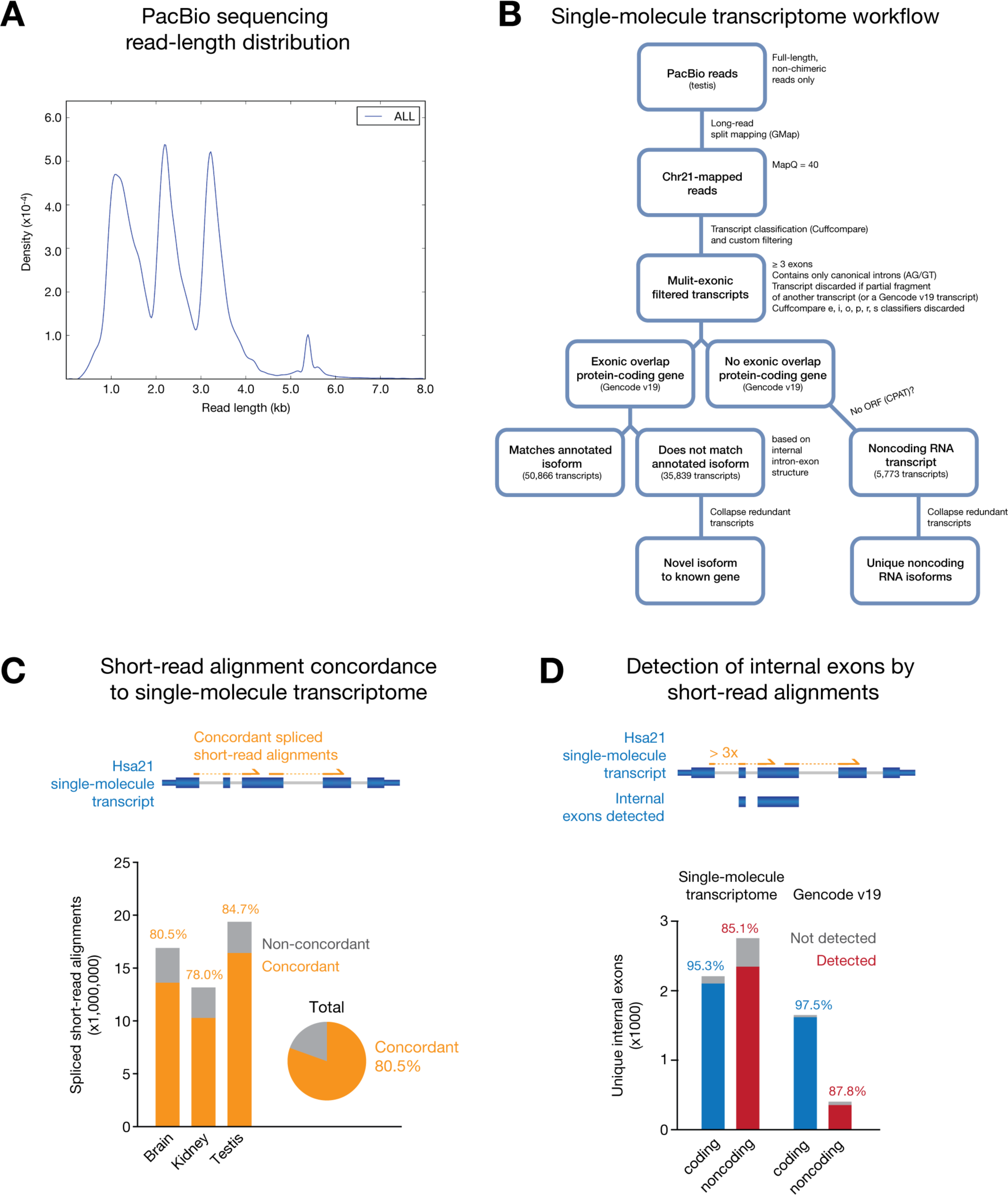
Single-molecule Hsa21 transcriptome workflow and statistics. Deep single-molecule PacBio sequencing was performed on Hsa21-enriched cDNA from human testis. **(A)** Length distributions for full-length non-chimeric (FLNC) sequencing reads in PacBio library. Peaks and valleys reflect outcomes of fragment size selection (**Materials & Methods**). **(B)** Flow-chart summarizes the process by which PacBio FLNC reads were mapped, filtered and classified to generate a single-molecule Has21 transcriptome profile (**Materials & Methods**). **(C)** Concordance of spliced short-read alignment junctions to introns in the Hsa21 single-molecule transcriptome. Proportions of total spliced alignments are shown for each tissue (bars) and combined tissues libraries (pie chart). **(D)** Detection of coding/noncoding unique internal exons from Hsa21 single-molecule transcriptome by spliced short-read alignments (both boundaries specified by ≥ 3 spliced short-read alignment termini) compared to unique coding/noncoding introns in the Gencode (v19) catalog.

**Figure S4. Related to Figure 1.**
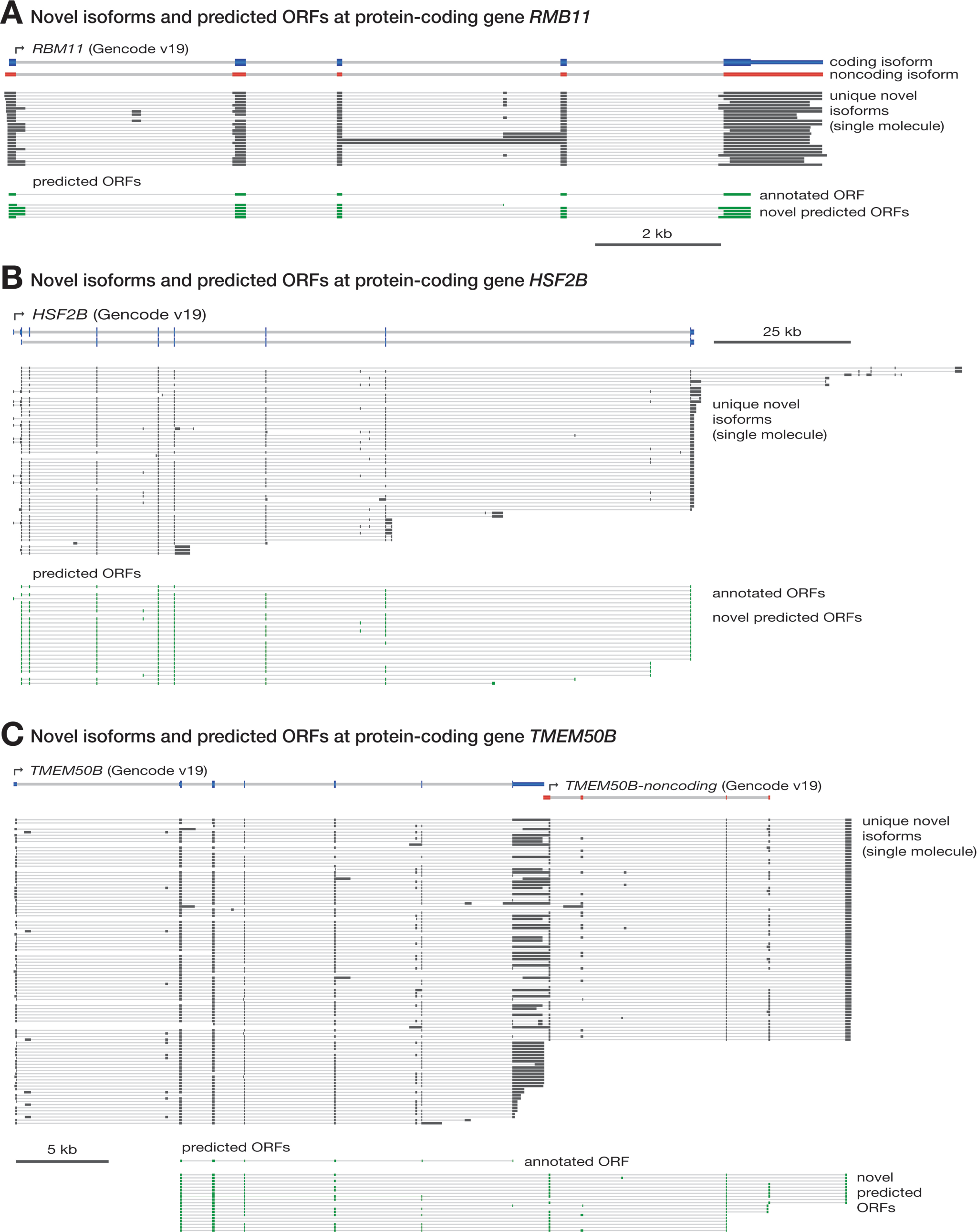
Examples of novel isoforms and predicted open reading frames on Hsa21. Singlemolecule RNA CaptureSeq identified 7,310 unique multi-exonic isoforms (including noncoding isoforms) at protein-coding loci on Hsa21, of which 77% were novel. These encoded up to 2,365 distinct, non-annotated predicted open reading frames (ORFs). Genome browser views show examples of annotated isoforms (Gencode v19), unique singlemolecule isoforms and novel predicted ORFs at three protein-coding genes **(A)** *RBM11*, **(B)** *HSF2B* and **(C)** *TMEM50B*. For *TMEM50B*, an adjacent lncRNA is incorporated by splicing to produce several novel extended ORFs (in frame with the annotated ORF for *TMEM50B).*

**Figure S5. Related to Figure 1.**
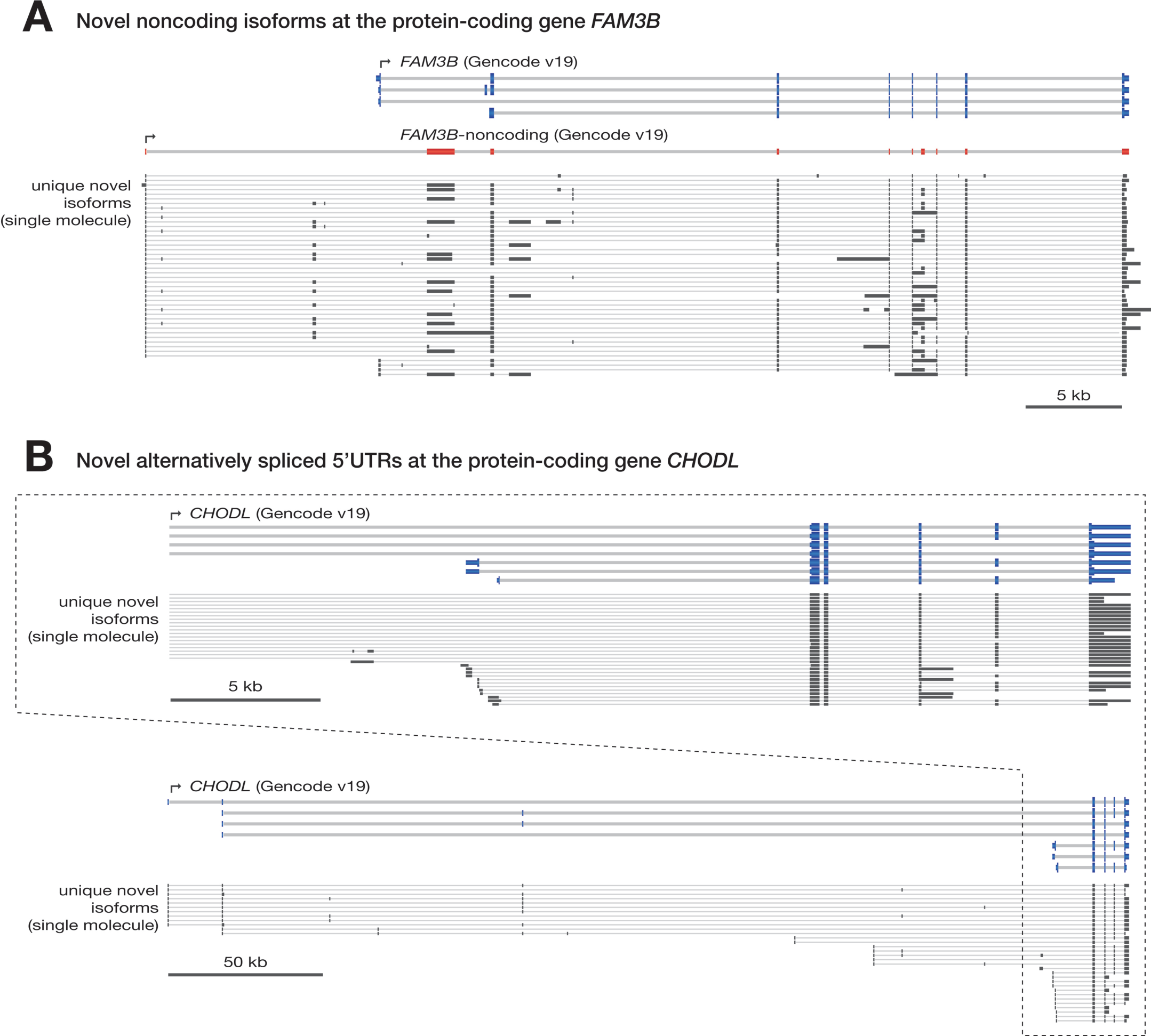
Examples of novel noncoding isoforms and alternatively spliced UTRs on Hsa21. The majority of novel isoforms to protein-coding genes detected using single-molecule RNA CaptureSeq were noncoding variants or possessed novel untranslated region (UTR) variants, rather than novel coding sequences. Alternative splicing of UTRs (both 5′ and 3′) was common and often highly complex. Genome browser views show examples of annotated isoforms (Gencode v19) and unique single-molecule isoforms at **(A)** *FAM3B* and **(B)** *CHODL. FAM3B* encodes numerous novel noncoding isoforms, in addition to a single annotated noncoding isoform. *CHODL* encodes a long, spliced 5′UTR, which includes multiple previously non-annotated exons. In both cases, noncoding exons (both UTR exons and exons exclusive to noncoding isoforms) are alternatively spliced, whereas exons inside open reading frames are typically constitutive. This manifests in a large diversity of novel noncoding isoforms and alternative UTR variants.

**Figure S6. Related to Figure 1.**
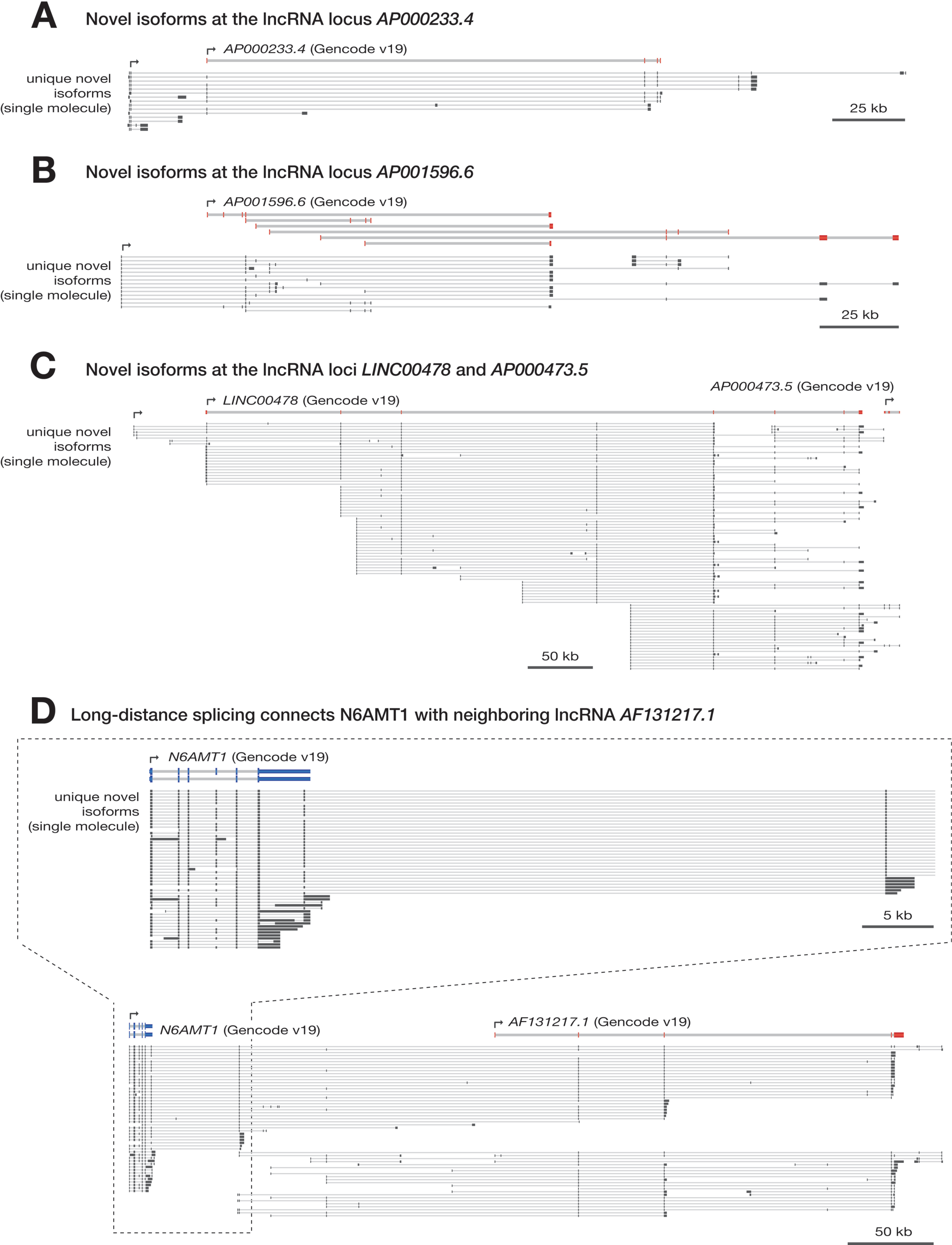
Examples of novel highly spliced lncRNA isoforms on Hsa21. Our analysis identified 1,589 novel lncRNA transcripts across Hsa21, encompassing 1,663 unique internal exons and 3,210 unique canonical introns. More complete gene models were generated for many known lncRNAs. Multiple partial lncRNA annotations were sometimes incorporated into a single unified loci and, in several cases, lncRNAs were spliced with neighboring protein-coding genes to form long, non-annotated, UTRs. Genome browser views show examples of unique singlemolecule isoforms for known and novel lncRNAs, including **(A)** *AP000233.4,* **(B)** *AP001596.6*, **(C)** *LINC00478* and *AP0004735,* which are linked by splicing, and **(D)** *AF131217.1,* which forms a novel 3′UTR to the upstream protein-coding gene *N6AMT1.* In all examples, noncoding exons are typically alternatively spliced, manifesting in a large diversity of novel lncRNA isoforms.

**Figure S7. Related to Figure 2.**
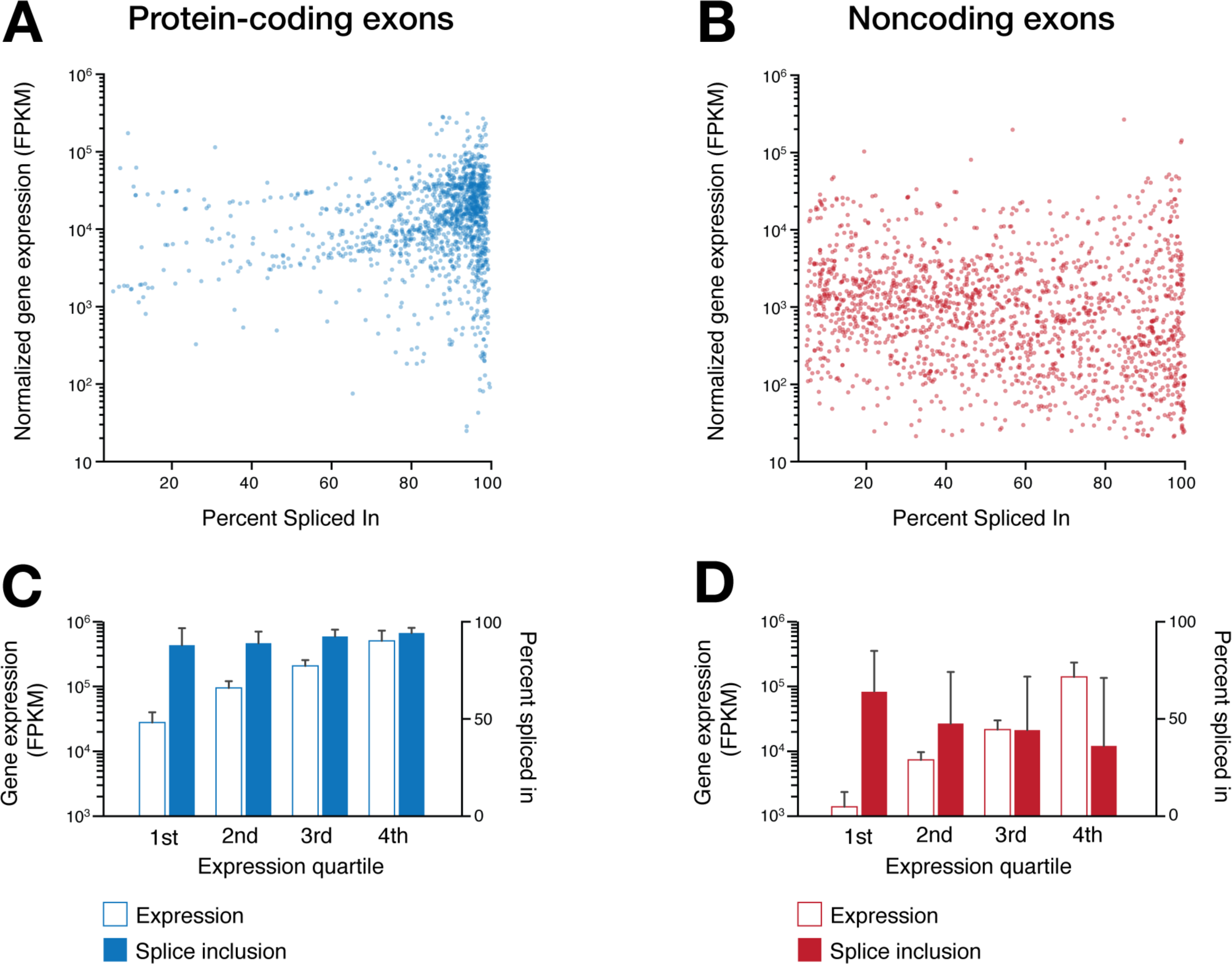
Gene expression and Percent Splice Inclusion are independent metrics. To assess alternative splicing we calculated Percent Splice Inclusion scores for all internal exons in our Hsa21 transcriptome (PSI; **Materials & Methods**). We observed a strong depletion of PSI scores among noncoding exons (median PSI = 55.5%) compared to protein-coding exons (median PSI = 90.5%; **Figure 2B**), implying enriched alternative splicing. To confirm that this was not simply an artifact of their low abundance, we explored the relationship between PSI scores and gene expression. **(A-D)** The relationship between overall gene expression (FPKM) and PSI scores for protein-coding **(A)** and noncoding **(B)** exons. Bar charts below summarize gene expression (FPKM) and PSI scores for protein-coding **(C)** and noncoding **(D)** exons within each gene expression quartile (mean ± SD). These analyses show that exon-PSI and overall gene expression metrics are independent from one and other.

**Figure S8. Related to Figure 2.**
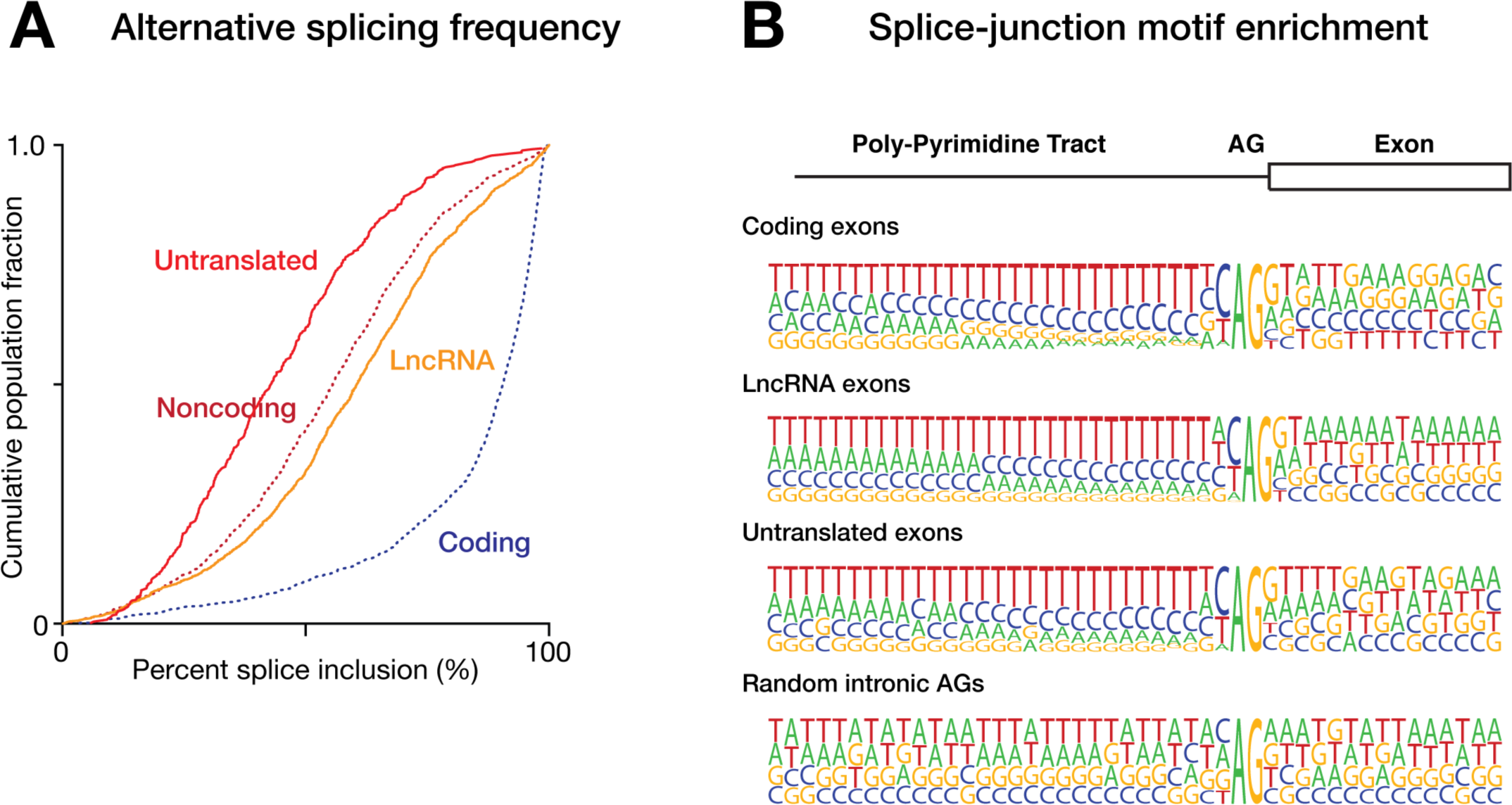
Noncoding exons exhibit enriched alternative splicing and possess *bona fide* splice sites. **(A)** Cumulative frequency distributions show percent splice inclusion (PSI) scores for protein-coding and noncoding internal exons in human tissues. The noncoding exon population is further parsed into lncRNA exons and noncoding exons belonging to protein-coding loci. **(B)** Nucleotide frequency plots illustrate canonical poly-pyrimidine enrichment immediately upstream of splice acceptor sites for protein-coding and noncoding exons from both lncRNAs and protein-coding loci. Randomly selected intronic AG dinucleotides are shown for comparison and, as expected, exhibit no poly-pyrimidine enrichment.

**Figure S9. Related to Figure 2.**
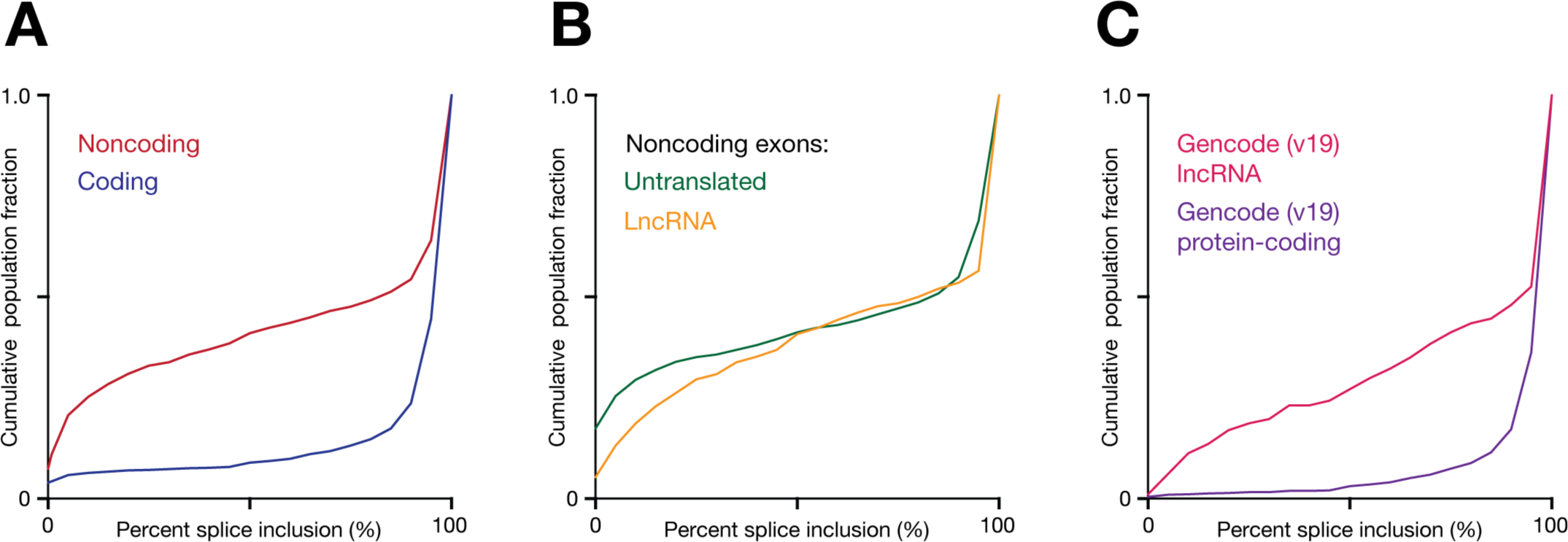
Percent splicing inclusions (PSI) scores calculated with PacBio sequencing reads. The PSI scores presented in **Figure 2B** were calculated by counting spliced short-read alignments that include/exclude exons represented in our single-molecule Has21 transcriptome (**Materials and Methods**). As an alternative to this combined approach, PSI scores were independently calculated using only spliced PacBio alignments (no short-reads). **(A-C)** Cumulative frequency distributions show PSI scores for unique internal exons calculated using only spliced PacBio alignments. Comparison are: **(A)** all protein-coding and noncoding exons detected on Hsa21; **(B)** noncoding exons, further parsed into lncRNA and untranslated exons at protein-coding loci; **(C)** exons from Gencode (v19) protein-coding genes and lncRNAs.

**Figure S10. Related to Figure 2.**
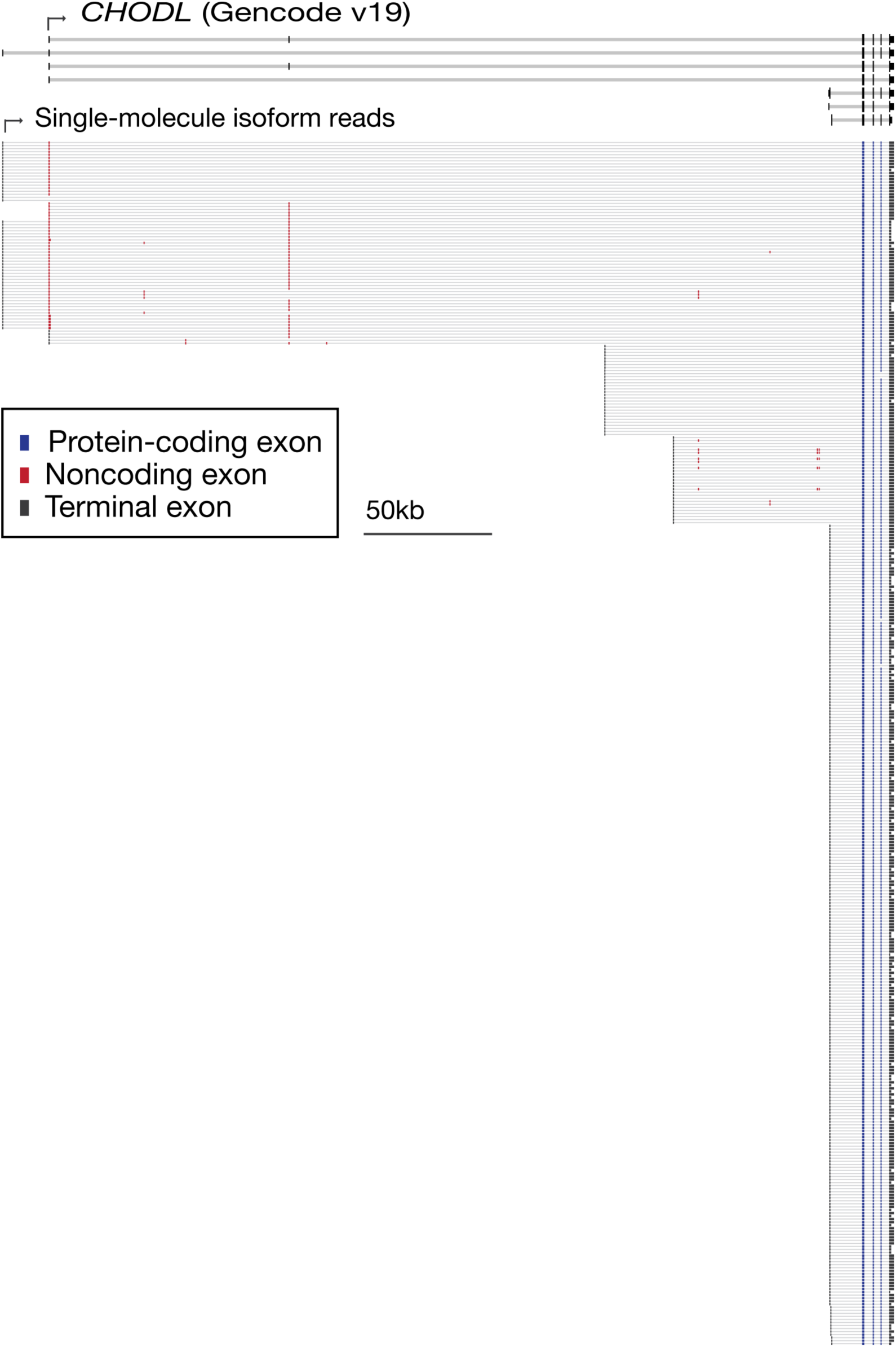
Example of enriched alternative splicing among untranslated exons. Annotated transcripts (Gencode v19) and mapped single-molecule isoform reads at the protein-coding gene *CHODL. CHODL* provides an illustrative example of enriched alternative splicing among UTR exons (in this case within the 5'UTR), relative to protein-coding exons at the same locus. Internal exons are identified as protein-coding (blue) or noncoding (red).

**Figure S11. Related to Figure 2.**
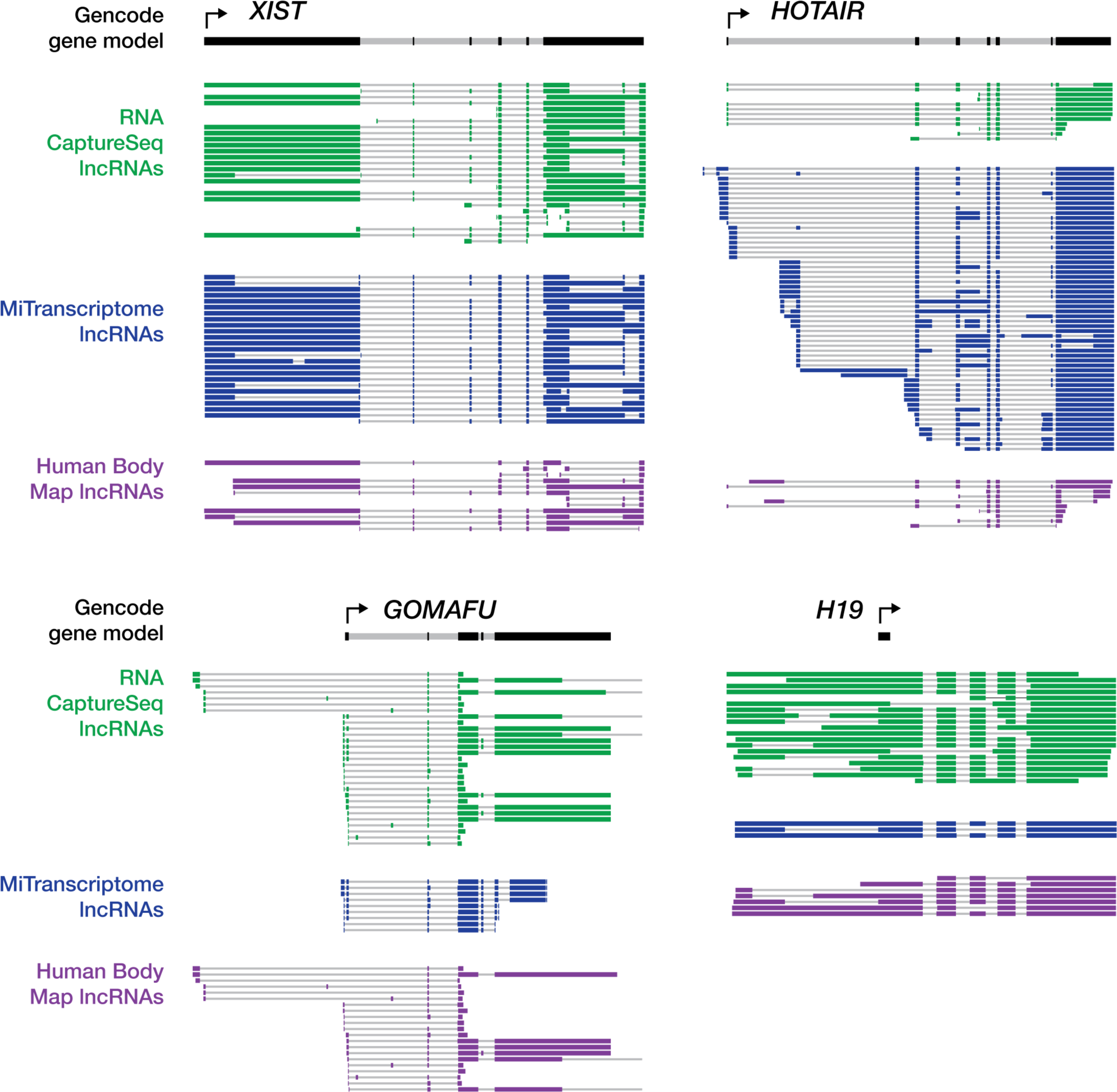
Universal alternative splicing among well-characterized functional lncRNAs. We obtained transcriptome annotations developed by (green; Clark et al., 2015), who applied RNA CaptureSeq on pooled human tissues to profile lncRNA loci at high depth, as well as the MiTranscriptome (blue; Iyer et al., 2015) and Human Body Map (purple; Cabili et al., 2011) lncRNA catalogs. With these data we scrutinized four well-known, functionally characterized lncRNAs – *XIST*, *HOTAIR*, *GOMAFU* and *H19* – for evidence of universal alternative splicing. All four of these lncRNAs exhibit complex alternative splicing patterns and, strikingly, almost no internal splice site in any of these examples is constitutively spliced across the three datasets.

**Figure S12. Related to Figure 3.**
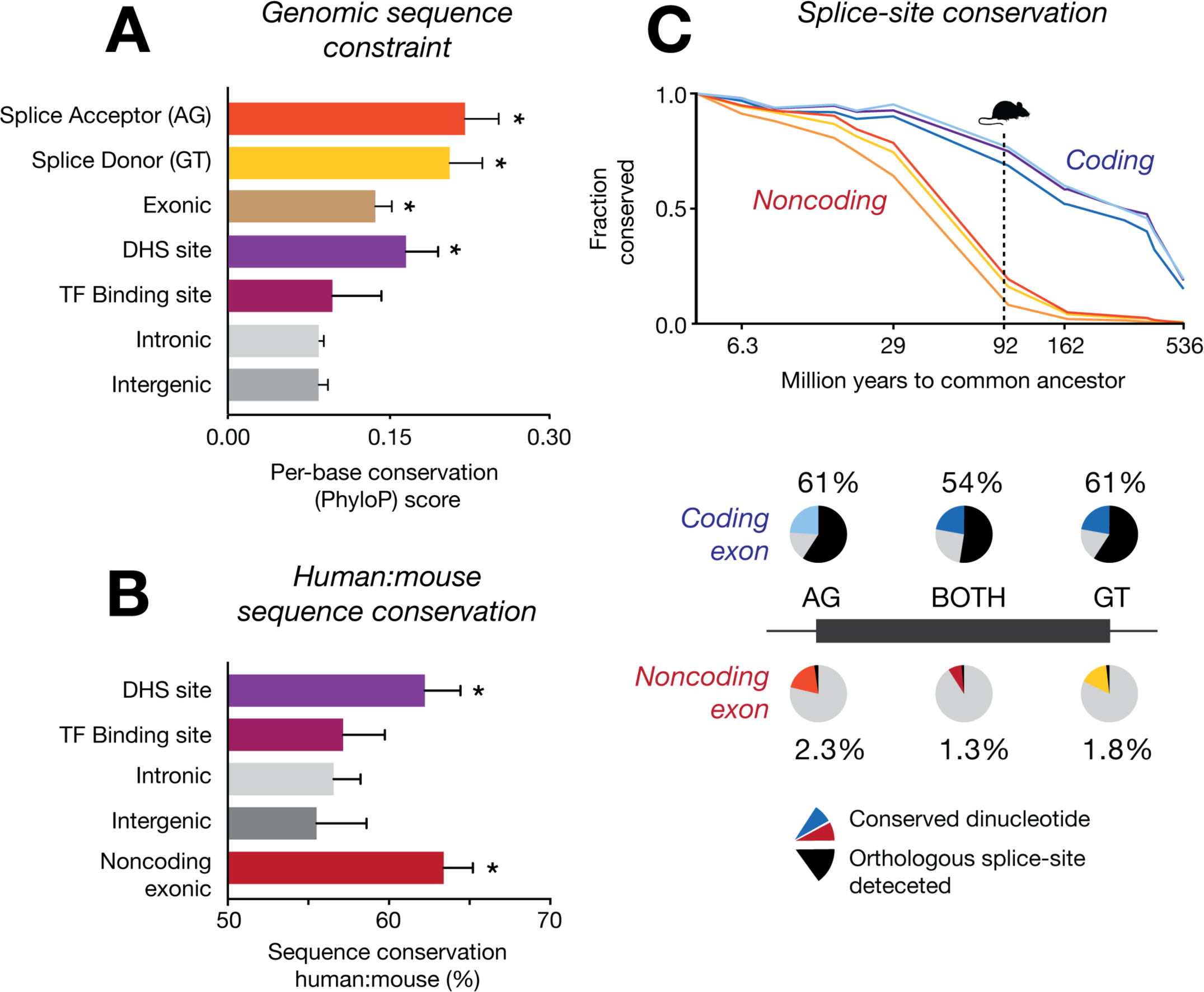
Evolutionary conservation of noncoding exons on Hsa21. **(A)** Per-base conservation scores (PhyloP scores from 100 Vertebrate MultiZ alignments) for the following features on Hsa21: novel noncoding exons, noncoding splice site dinucleotides (AG/GT), DNase Hypersensitive (DHS) sites, Transcription Factor (TF) binding sites and randomly selected intronic and intergenic sequences (mean ± SD, asterisks denote significant increases relative to intergenic sequences, p < 0.0001, unpaired t-test). **(B)** Percentage human:mouse sequence similarity among all alignable features for noncoding exons, DHS sites, TF binding sites and randomly selected intergenic or intronic sequences. **(C;** upper) The fraction of coding/noncoding splice-site dinucleotides (AG/GT/both) detected on Hsa21 that are conserved in other vertebrate genomes (arranged by Million years to common ancestor). **(C;** lower) Pie-charts indicate the proportion of Hsa21 exons with splice-site dinucleotides that are conserved in the mouse genome (red/blue) and the proportion for which an equivalent splice site could also be detected in mouse RNA CaptureSeq libraries (black).

**Figure S13 (previous page). Related to Figure 3, 4.**
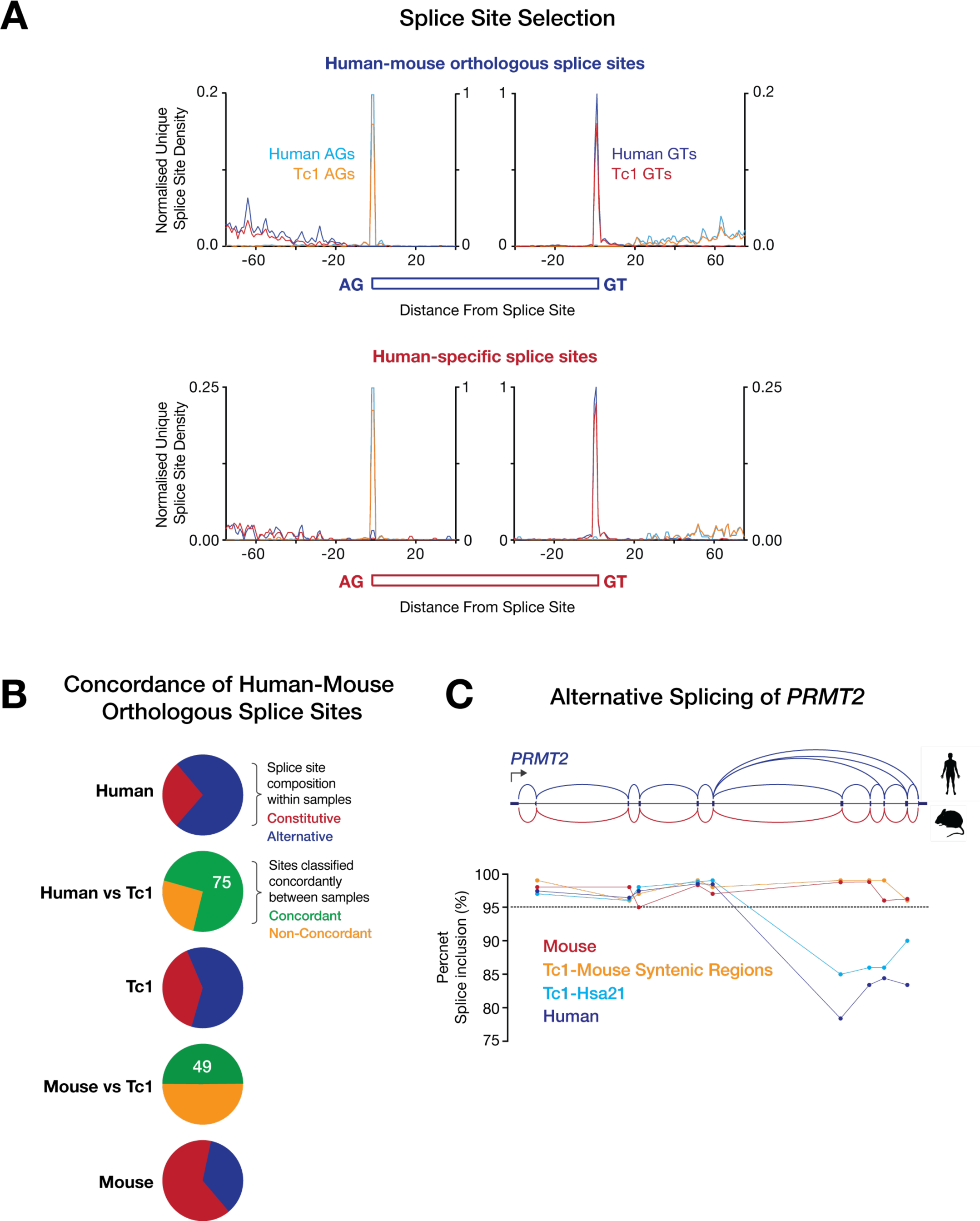
Splicing concordance of splice sites between human and Tc1 mouse libraries. **(A)** Density plot indicates position of unique splice sites identified on Hsa21 in Tc1 mouse libraries (brain, *n* = 3) relative to unique splice sites identified in matched human libraries (*n* = 2). Splice sites that have orthologs present between human and mouse are indicated in upper panel, whilst human-specific splice sites that distinguish human genes from their mouse orthologs are indicated in lower panel. The analysis suggests that human splice site selection is faithfully recapitulated in the Tc1 mouse. **(B)** All orthologous splice sites were classified as constitutive (PSI > 95) or alternative in human and mouse samples, as well as those deriving from Hsa21 in the Tc1 mouse. Pie charts (red/blue) indicate the proportion that were classified as constitutive/alternative in each sample, while intervening charts (green/orange) depict the proportion of sites that were concordantly classified between samples. The analysis shows that exon inclusion on Hsa21 in the Tc1 mouse resembles human, rather than mouse samples **(C)** This is illustrated by splicing profiles for the conserved *PRMT2* gene. Plot indicates the PSI scores for each exon assembled in the *PRMT2* locus in human, mouse and Tc1 libraries, deriving either from Hsa21 or mouse syntenic regions. Human-specific alternative splicing of the *PRMT2* gene (encoded on Hsa21) is observed in the Tc1 mouse (Tc1-Hsa21). By contrast, splicing of all exons is constitutive at the *Prmt2* locus (encoded in the mouse genome) in Tc1 and wild type mouse samples.

**Figure S14. Related to Figure 4.**
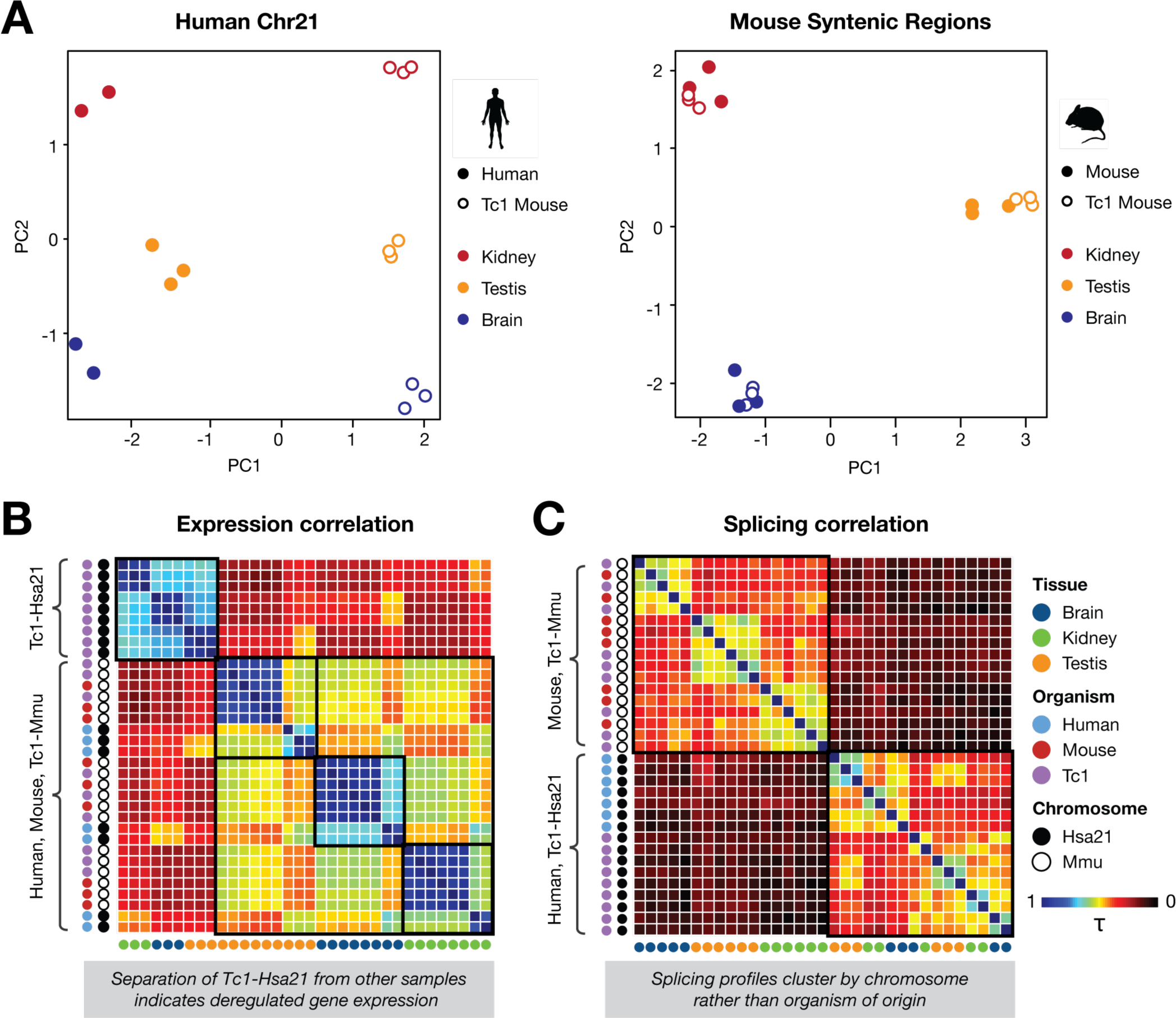
Comparison of gene expression and splicing in human, mouse and Tc1 mouse tissues. **(A)** Principle component analysis of gene expression for human-mouse orthologous gene pairs on Hsa21 in human and Tc1 tissue libraries and within mouse syntenic regions for mouse and Tc1 tissue libraries. PCA plots show human gene expression (from Hsa21) is broadly deregulated in Tc1, whilst mouse gene expression (from syntenic regions in the mouse genome) is unperturbed in Tc1 tissues. **(B)** Correlation matrices (based on Kendall rank correlation coefficient; τ) of gene expression for orthologous gene pairs in human, mouse and Tc1 tissue samples. Black boxes indicate grouping of human, mouse and Tc1 samples according to tissue, with the exception of human genes in Tc1 mouse samples, which clustered separately, implying deregulated expression. **(C)** Correlation matrices (τ) of percentage splice inclusion scores in human, mouse and Tc1 tissue samples shows the grouping of samples according to chromosome (as indicated by black boxes), rather than tissue or organism source.

**Figure S15. Related to Figure 4.**
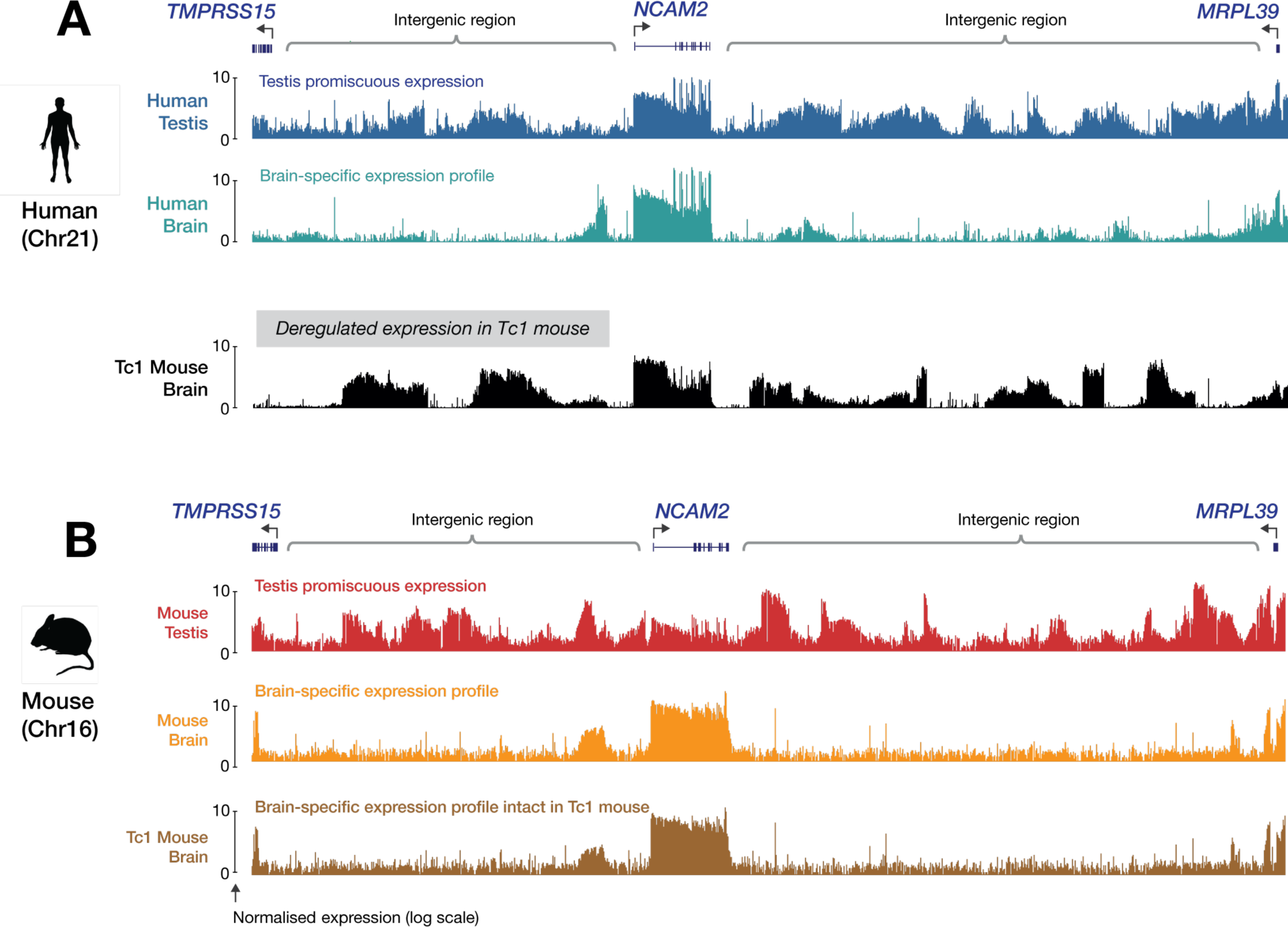
Human and mouse *NCAM2* loci and flanking intergenic regions in human, mouse and Tc1 mouse. Plots indicate transcriptional activity (log scale) recorded at the *NCAM2* locus and its flanking intergenic regions on **(A)** Hsa21 and **(B)** the syntenic region of the mouse genome, in human, mouse and Tc1 mouse tissues. Notably, the brain-specific expression/silencing profile of lncRNAs observed in human (Hsa21; teal) was deregulated in the Tc1 mouse brain (Hsa21; black), with lncRNAs being promiscuously expressed. By contrast, the mouse brain expression profile (mouse syntenic regions; orange) was maintained in Tc1 mouse (mouse syntenic regions; brown), indicating deregulated expression in Tc1 was restricted to the human chromosome.

**Supplementary Table 1.Related to Figure 1.** Library statistics for short-read RNA-Seq and CaptureSeq analysis of human K562 cells.

| Library | Organism | Material | Method | Raw reads<br>(millions) | Unique alignments (hg19; millions) |  |  | Capture<br>enrichment |
| --- | --- | --- | --- | --- | --- | --- | --- | --- |
|  |  |  |  |  | Off-target | On-target | On-rate |  |
| K562_rnaseq_1 | Human | K562 | RNA-Seq | 145.90 | 114.54 | 1.46 | 0.013 |  |
| K562_rnaseq_2 | Human | K562 | RNA-Seq | 151.52 | 116.95 | 1.46 | 0.012 |  |
| K562_rnaseq_3 | Human | K562 | RNA-Seq | 126.94 | 97.30 | 1.47 | 0.015 |  |
| K562_capture_1 | Human | K562 | CaptureSeq | 158.84 | 33.77 | 86.04 | 0.718 | 54-fold |
| K562_capture_2 | Human | K562 | CaptureSeq | 170.02 | 37.50 | 96.41 | 0.720 |  |
| K562_capture_3 | Human | K562 | CaptureSeq | 147.66 | 30.03 | 75.66 | 0.716 |  |

**Supplementary Table 2. Related to Figure 1.**
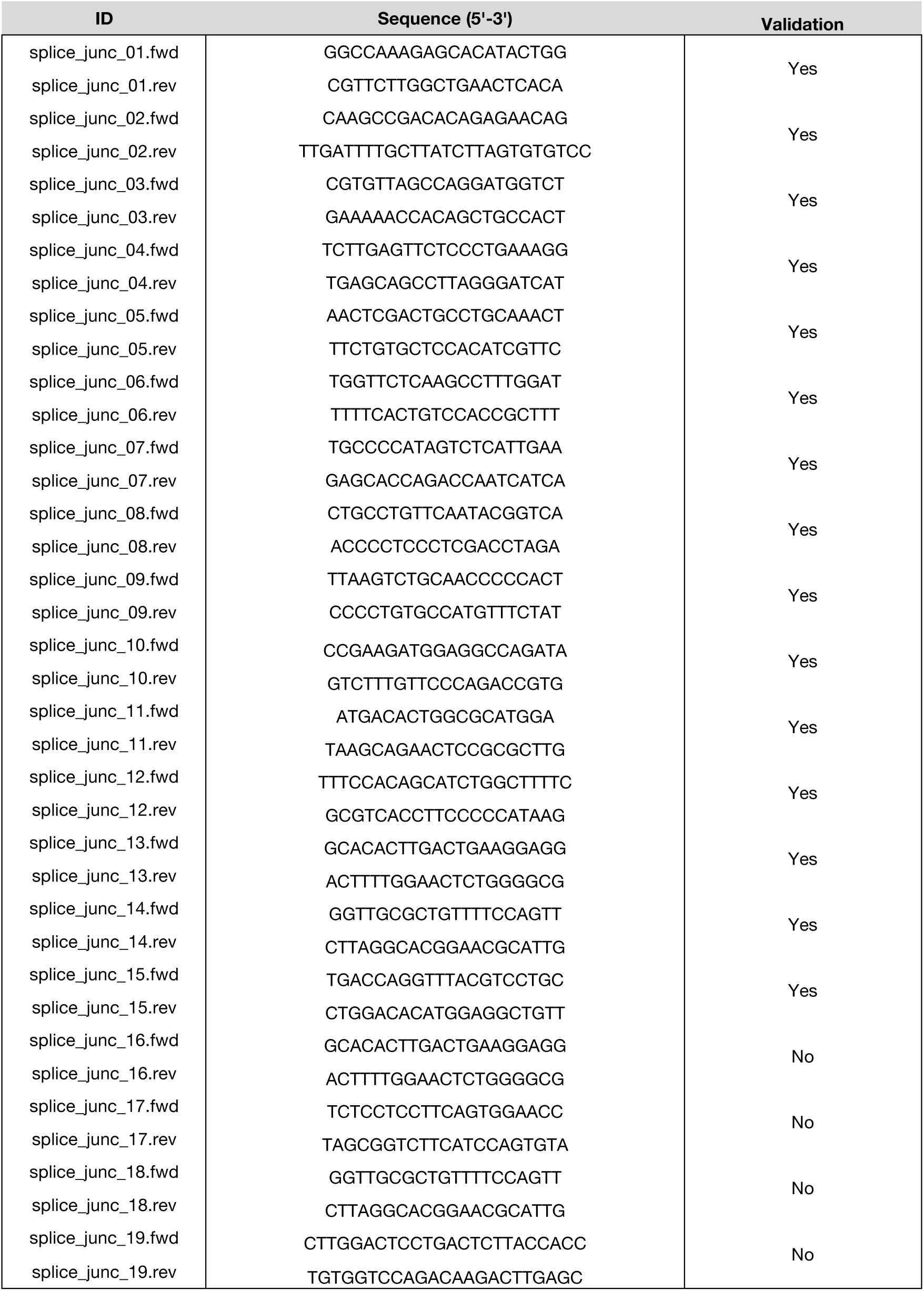
Primer sequences for validation of novel splice junctions by RT-PCR and Sanger sequencing.

**Supplementary Table 3. Related to Figure 1.** Library statistics for PacBio single-molecule RNA CaptureSeq analysis of human testis

|  |  |
| --- | --- |
| **Library** | hsa_testis_pacbio |
| **Organism** | Human |
| **Material** | Testis |
| **Method** | Hsa21-Capture, PacBio IsoSeq |
| **SMRT Cells** | 24 |
| **Circular consensus sequences** | 2,011,066 |
| **Full-length non-chimeric (FNLC) reads** | 843,326 |
| **FLNC reads aligned to Hsa21** | 387,029 |
| **On-target rate** | 0.46 |
| **Capture enrichment** | 35-fold |
| **Average FLNC length** | 2.35 kb |
| **Total FNLC** | 910 Mb |
| **Full-length multi-exonic transcripts** | 101,478 |

**Supplementary Table 4. Related to Figure 1.** Library statistics for short-read RNA CaptureSeq analysis of human tissues.

| Library | Organism | Material | Method | Raw reads<br>(millions) | Unique alignments (hg19; millions) |  |  | Capture<br>enrichment |
| --- | --- | --- | --- | --- | --- | --- | --- | --- |
|  |  |  |  |  | Off-target | On-target | On-rate |  |
| hsa_brain_1 | Human | Brain | CaptureSeq | 67.02 | 10.14 | 52.01 | 0.837 | 65-fold |
| hsa_brain_2 | Human | Brain | CaptureSeq | 119.32 | 5.66 | 50.44 | 0.899 |  |
| hsa_kidney_1 | Human | Kidney | CaptureSeq | 68.56 | 10.23 | 52.76 | 0.838 | 64-fold |
| hsa_kidney_2 | Human | Kidney | CaptureSeq | 244.66 | 13.54 | 97.94 | 0.879 |  |
| hsa_testis_1 | Human | Testis | CaptureSeq | 144.90 | 13.51 | 87.53 | 0.866 | 66-fold |
| hsa_testis_2 | Human | Testis | CaptureSeq | 111.42 | 8.09 | 68.26 | 0.894 |  |
| hsa_testis_3 | Human | Testis | CaptureSeq | 143.64 | 6.59 | 48.38 | 0.880 |  |

**Supplementary Table 5. Related to Figure 1.** Comparison of Hsa21 single-molecule transcriptome to existing catalogs. Total number of isoforms, unique internal exons and unique canonical introns identified on Hsa21 and the number recovered from the Gencode (v19), MiTranscriptome (v2) and FANTOM CAT (lv2) catalogs, as well as the percentage of CaptureSeq transcripts/exons/introns that are novel w.r.t. all three annotations.

|  | CaptureSeq | Number recovered from existing annotations |  |  |  | Novel (%) |
| --- | --- | --- | --- | --- | --- | --- |
|  |  | Gencode | MiT | FANTOM | Combined |  |
| **Total unique:** |  |  |  |  |  |  |
| Isoforms | 8908 | 1494 | 1078 | 1846 | 2178 | 76% |
| Exons | 4913 | 2443 | 2679 | 2847 | 3146 | 36% |
| Introns | 9575 | 3806 | 4117 | 4759 | 5171 | 46% |
| **Protein-coding loci** |  |  |  |  |  |  |
| Isoforms | 7310 | 1116 | 718 | 1410 | 1663 | 77% |
| Coding exons | 2185 | 1788 | 1730 | 1840 | 1894 | 13% |
| Coding introns | 3639 | 2396 | 2222 | 2646 | 2794 | 23% |
| Noncoding exons<br>(untranslated) | 1746 | 355 | 443 | 600 | 653 | 63% |
| Noncoding introns<br>(untranslated) | 2726 | 478 | 616 | 847 | 921 | 66% |
| **Noncoding loci** |  |  |  |  |  |  |
| Isoforms | 1598 | 378 | 360 | 436 | 515 | 68% |
| Exons | 1663 | 476 | 742 | 658 | 911 | 45% |
| Introns | 3210 | 931 | 1278 | 1266 | 1456 | 55% |

**Supplementary Table 6. Related to Figure 1.** Transcript model attributes for Hsa21 single-molecule transcriptome. Median sizes for isoforms, unique internal exons and unique canonical introns within annotated (Gencode v19) or CaptureSeq transcripts on Hsa21.

|  | Median size (bp) |  |
| --- | --- | --- |
|  | Gencode<br>(v19) | CaptureSeq |
| **Total unique:** |  |  |
| Isoforms | 1404 | 2218 |
| Exons | 125 | 130 |
| Introns | 2371 | 2761 |
| **Protein-coding loci** |  |  |
| Isoforms | 1077 | 2322 |
| Coding exons | 124 | 122 |
| Coding introns | 2257 | 2300 |
| Noncoding exons<br>(untranslated) | 136 | 158 |
| Noncoding introns<br>(untranslated) | 2589 | 2883 |
| **Noncoding loci** |  |  |
| Isoforms | 700 | 1582 |
| Exons | 116 | 123 |
| Introns | 2956 | 3847 |

**Supplementary Table 7. Related to Figure 3.** Library statistics for short-read RNA CaptureSeq analysis of mouse tissues.

| Library | Organism | Material | Method | Raw reads<br>(millions) | Unique alignments (mm10; millions) |  |  | Capture<br>enrichment |
| --- | --- | --- | --- | --- | --- | --- | --- | --- |
|  |  |  |  |  | Off-target | On-target | On-rate |  |
| mmu_brain_1 | WT mouse | Brain | CaptureSeq | 79.00 | 7.11 | 66.37 | 0.903 | 64-fold |
| mmu_brain_2 | WT mouse | Brain | CaptureSeq | 117.82 | 5.57 | 94.21 | 0.944 |  |
| mmu_brain_3 | WT mouse | Brain | CaptureSeq | 133.66 | 29.80 | 75.28 | 0.716 |  |
| mmu_kidney_1 | WT mouse | Kidney | CaptureSeq | 105.10 | 26.47 | 47.90 | 0.644 | 58-fold |
| mmu_kidney_2 | WT mouse | Kidney | CaptureSeq | 132.68 | 5.25 | 114.08 | 0.956 |  |
| mmu_kidney_3 | WT mouse | Kidney | CaptureSeq | 184.54 | 41.95 | 104.99 | 0.715 |  |
| mmu_testis_1 | WT mouse | Testis | CaptureSeq | 102.14 | 4.91 | 86.80 | 0.946 | 68-fold |
| mmu_testis_2 | WT mouse | Testis | CaptureSeq | 107.24 | 3.48 | 98.44 | 0.966 |  |
| mmu_testis_3 | WT mouse | Testis | CaptureSeq | 47.28 | 7.61 | 32.76 | 0.811 |  |

**Supplementary Table 8. Related to Figure 3,4.**
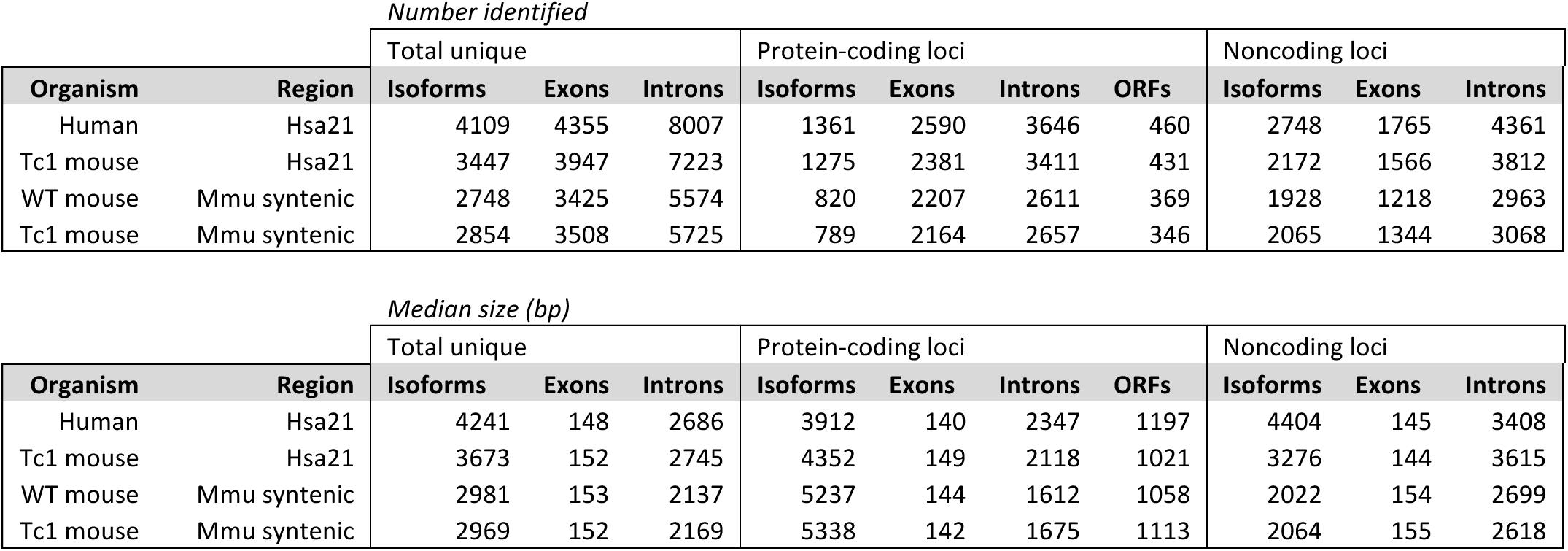
Summary statistics for human, mouse and Tc1 mouse transcriptome assemblies. Total number of isoforms, unique internal exons and unique canonical introns identified on Hsa21 and/or mouse syntenic regions and median sizes for each feature.

**Supplementary Table 9. Related to Figure 3,4.** Library statistics for RNA CaptureSeq analysis of Tc1 mouse tissues.

| Library | Organism | Material | Method | Raw reads<br>(millions) | Unique alignments (hg19; millions) |  |  | Capture<br>enrichment |
| --- | --- | --- | --- | --- | --- | --- | --- | --- |
|  |  |  |  |  | Off-target | On-target | On-rate |  |
| tc1_brain_1 | Tc1 mouse | Brain | CaptureSeq | 144.40 | 1.59 | 70.10 | 0.978 | 74-fold |
| tc1_brain_2 | Tc1 mouse | Brain | CaptureSeq | 84.38 | 0.54 | 27.08 | 0.980 |  |
| tc1_brain_3 | Tc1 mouse | Brain | CaptureSeq | 139.00 | 0.90 | 37.54 | 0.977 |  |
| tc1_kidney_1 | Tc1 mouse | Kidney | CaptureSeq | 103.30 | 0.83 | 37.42 | 0.978 | 74-fold |
| tc1_kidney_2 | Tc1 mouse | Kidney | CaptureSeq | 143.98 | 0.62 | 22.21 | 0.973 |  |
| tc1_kidney_3 | Tc1 mouse | Kidney | CaptureSeq | 137.36 | 0.41 | 29.89 | 0.986 |  |
| tc1_testis_1 | Tc1 mouse | Testis | CaptureSeq | 125.38 | 0.82 | 41.28 | 0.980 | 72-fold |
| tc1_testis_2 | Tc1 mouse | Testis | CaptureSeq | 138.64 | 0.71 | 40.08 | 0.982 |  |
| tc1_testis_3 | Tc1 mouse | Testis | CaptureSeq | 132.22 | 0.82 | 53.12 | 0.985 |  |

| Library | Organism | Material | Method | Raw reads<br>(millions) | Unique alignments (mm10; millions) |  |  | Capture<br>enrichment |
| --- | --- | --- | --- | --- | --- | --- | --- | --- |
|  |  |  |  |  | Off-target | On-target | On-rate |  |
| tc1_brain_1 | Tc1 mouse | Brain | CaptureSeq | 144.40 | 8.99 | 61.19 | 0.872 | 61-fold |
| tc1_brain_2 | Tc1 mouse | Brain | CaptureSeq | 84.38 | 4.76 | 48.93 | 0.911 |  |
| tc1_brain_3 | Tc1 mouse | Brain | CaptureSeq | 139.00 | 24.93 | 46.08 | 0.649 |  |
| tc1_kidney_1 | Tc1 mouse | Kidney | CaptureSeq | 103.30 | 6.15 | 55.62 | 0.900 | 58-fold |
| tc1_kidney_2 | Tc1 mouse | Kidney | CaptureSeq | 143.98 | 29.55 | 58.49 | 0.664 |  |
| tc1_kidney_3 | Tc1 mouse | Kidney | CaptureSeq | 137.36 | 24.02 | 65.04 | 0.730 |  |
| tc1_testis_1 | Tc1 mouse | Testis | CaptureSeq | 125.38 | 4.39 | 72.33 | 0.943 | 66-fold |
| tc1_testis_2 | Tc1 mouse | Testis | CaptureSeq | 138.64 | 21.62 | 56.01 | 0.722 |  |
| tc1_testis_3 | Tc1 mouse | Testis | CaptureSeq | 132.22 | 4.32 | 71.30 | 0.943 |  |

**Supplementary Table 10. Related to Figure 1.** Performance statistics for each fragment-size fraction input for PacBio sequencing.

| Sample | FL % | # of FL ZMWs | Avg. FL len | FL length range | Artificial Chimeras |
| --- | --- | --- | --- | --- | --- |
| 0to2kb | 48 | 81,883 | 1,230 | 1,008-1,403 | 370 (0.22%) |
| 0to2kb_run2 | 64 | 164,549 | 1,328 | 1,108-1,521 | 1,547 (0.60%) |
| 1to3kb | 56 | 110,348 | 1,904 | 1,340-2,341 | 406 (0.21%) |
| 1to3kb_run2 | 50 | 204,445 | 2,235 | 2,072-2,477 | 1,507 (0.37%) |
| 3to6kb | 41 | 79,520 | 3,390 | 3,146-3,502 | 576 (0.30%) |
| 3to6kb_run2 | 42 | 164,348 | 3,437 | 3,170-3,542 | 1,132 (0.29%) |
| 4to10kb | 10 | 16,188 | 4,291 | 2,368-5,448 | 449 (0.27%) |
| 4to10kb_run2 | 10 | 22,045 | 4,681 | 3,458-5,459 | 446 (0.20%) |
| **TOTAL** |  | **843,326** |  |  |  |

